# A hardwired neural circuit for temporal difference learning

**DOI:** 10.1101/2025.09.18.677203

**Authors:** Malcolm G. Campbell, Yongsoo Ra, Zhiqin Chen, Shudi Xu, Mark Burrell, Sara Matias, Mitsuko Watabe-Uchida, Naoshige Uchida

**Affiliations:** Department of Molecular and Cellular Biology, Harvard University, Cambridge, MA, USA; Center for Brain Science, Harvard University, Cambridge, MA, USA; Program in Neuroscience, Harvard University, Boston, MA, USA; Department of Neurobiology, Northwestern University, Evanston, IL, USA

## Abstract

The neurotransmitter dopamine plays a major role in learning by acting as a teaching signal to update the brain’s predictions about rewards. A leading theory proposes that this process is analogous to a reinforcement learning algorithm called temporal difference (TD) learning, and that dopamine acts as the error term within the TD algorithm (TD error). Although many studies have demonstrated similarities between dopamine activity and TD errors^1–5^, the mechanistic basis for dopaminergic TD learning remains unknown. Here, we combined large-scale neural recordings with patterned optogenetic stimulation to examine whether and how the key steps in TD learning are accomplished by the circuitry connecting dopamine neurons and their targets. Replacing natural rewards with optogenetic stimulation of dopamine axons in the nucleus accumbens (NAc) in a classical conditioning task gradually generated TD error-like activity patterns in dopamine neurons by specifically modifying the task-related activity of NAc neurons expressing the D1 dopamine receptor (D1 neurons). In turn, patterned optogenetic stimulation of NAc D1 neurons in naïve animals drove dopamine neuron spiking according to the TD error of the stimulation pattern, indicating that TD computations are hardwired into this circuit. The transformation from D1 neurons to dopamine neurons could be described by a biphasic linear filter, with a rapid positive and delayed negative phase, that effectively computes a temporal difference. This finding suggests that the time horizon over which the TD algorithm operates—the temporal discount factor—is set by the balance of the positive and negative components of the linear filter, pointing to a circuit-level mechanism for temporal discounting. These results provide a new conceptual framework for understanding how the computations and parameters governing animal learning arise from neurobiological components.

## Introduction

The brain learns by comparing internal expectations against real outcomes and updating expectations when they are wrong. This idea—that a major function of the brain is to learn accurate predictions, and that it does so in part by computing and minimizing prediction errors—is an influential organizing principle in neuroscience^6^.

A prime example of this is the phasic activity of midbrain dopamine neurons, which canonically reflects the difference between predicted and actual reward—i.e. reward prediction error (RPE)—and acts as a teaching signal to update reward predictions (“value”)^1,3^. More specifically, phasic dopamine reflects a specific form of RPE called temporal difference RPE (TD RPE, or TD error), the error term in a reinforcement learning (RL) algorithm called temporal difference (TD) learning^7,8^ (**Fig. 1a**). TD learning is a cornerstone of most real-world successes of RL^8–11^.

**Figure 1:**
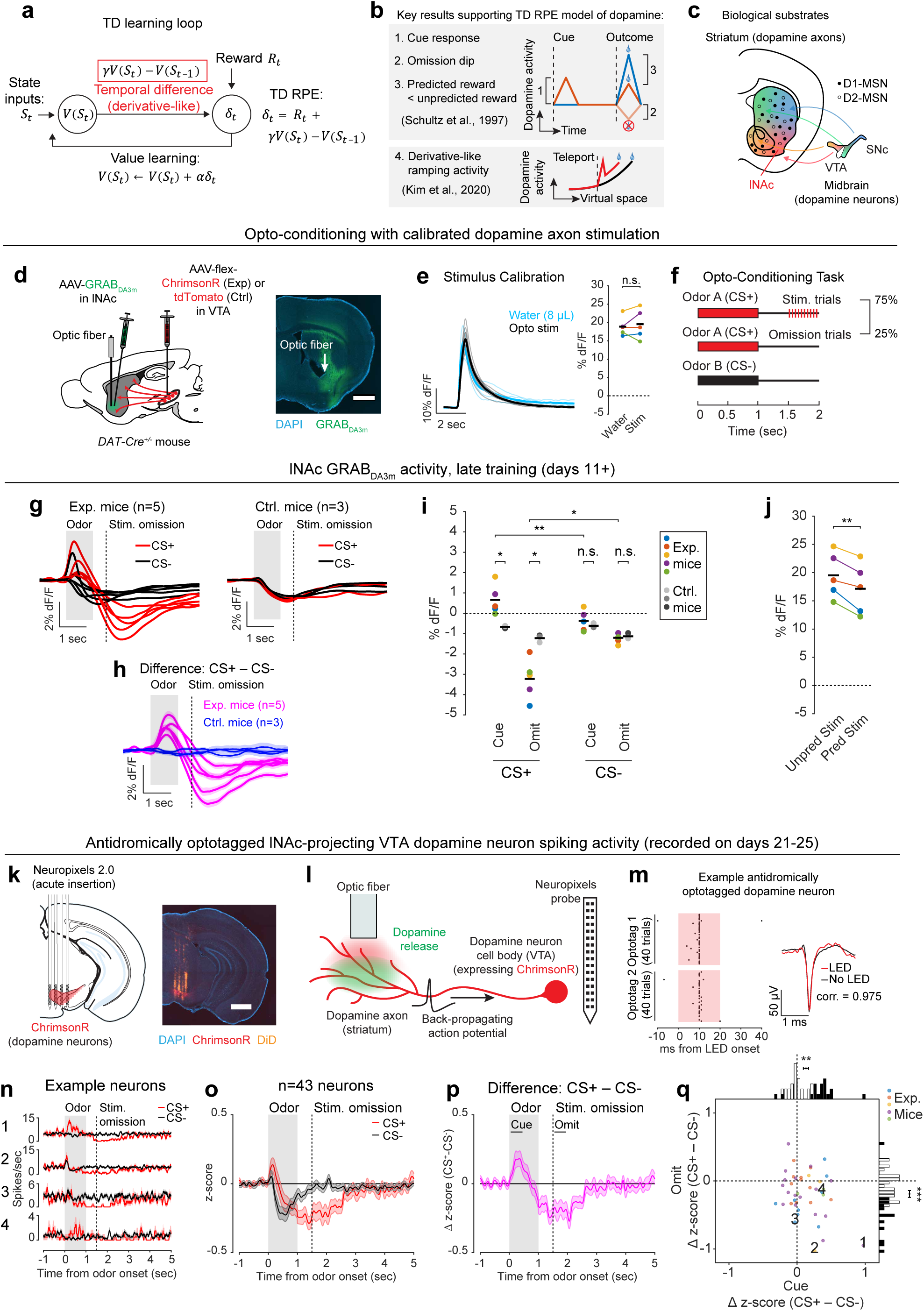
Opto-conditioning in lNAc generates TD error signals in dopamine activity. a) Schematic of a temporal difference (TD) learning loop^8^. The red box highlights the temporal difference calculation, which is exactly a derivative when the temporal discount factor *γ* = 1, but deviates from a derivative when *γ* ≠ 1. *S*_*t*_ : state at time *t*, *V*(*S*_*t*_): the value (expected discounted sum of future reward) associated with state *S*_*t*_, *R*_*t*_: reward at time *t*, *δ*_*t*_: TD RPE, *γ*: temporal discount factor, *α*: learning rate. b) Key results supporting the TD RPE model of dopamine. In addition to the classic results of Schultz et al. (1997)^1^ in classical conditioning tasks (cue response, omission dip, and expectation-dependent suppression of reward response), the derivative-like property of dopamine signaling in the absence of reward feedback was shown by Kim et al. (2020)^5^. c) TD learning is thought to involve bi-directional interactions between midbrain dopamine neurons and their primary output region, the striatum. We first focus on lateral nucleus accumbens (lNAc), a subregion of the striatum, because evidence suggests it plays a specialized role in value learning^32,38^. lNAc: lateral nucleus accumbens, VTA: ventral tegmental area, SNc: substantia nigra pars compacta, D1-MSN: D1 dopamine receptor-expressing medium spiny neuron, D2-MSN: D2 dopamine receptor-expressing medium spiny neuron. d) *Left:* Schematic of mouse surgery. ChrimsonR (exp. mice, *n* = 5) or tdTomato (ctrl. mice, *n* = 3) was expressed virally in VTA dopamine neurons and GRAB_DA3m_ in lNAc. The fiber targeting lNAc was implanted at an angle to allow acute Neuropixels penetrations targeting lNAc or VTA. *Right:* Example histology image showing fiber tip and GRAB_DA3m_ expression in lNAc. Scale bar = 1 mm. e) Stimulus calibration. Optogenetic stimulation parameters were chosen to match the dopamine response to a natural reward (8 *μ* water). *Left:* Each thin line represents the average GRAB_DA3m_ response to 8 *μ* water (cyan) or opto-stimulation (black) for an individual mouse (*n* = 5); thick lines show means over mice. *Right:* Average GRAB_DA3m_ signal in a 1-second window following water or stimulation delivery (mean %dF/F ± SEM, exp. mice [*n* = 5], water: 18.87 ± 1.16, stim: 19.52 ± 1.80, *P =* 0.56, paired t-test). Colored lines represent individual mice, black lines represent means over mice. n.s., Not Significant. f) Opto-conditioning task design. g) lNAc GRAB_DA3m_ photometry signals on omission trials, in late training (days 11+), in exp. mice (*n* = 5, left) or ctrl. mice (*n* = 3, right). Each pair of red/black lines represents the average GRAB_DA3m_ response to CS+ or CS-for an individual mouse, first averaged within day, then across days (days 11+). h) The average difference between CS+ and CS-GRAB_DA3m_ responses on omission trials, in late training (days 11+), for each mouse, colored by whether they expressed ChrimsonR (magenta) or tdTomato (blue) in dopamine neurons. Lines/shading show mean/SEM over late training sessions for each mouse. i) Average GRAB_DA3m_ signal during the odor cue (0-1 seconds after odor onset) or omission of opto-stimulation (1.5-2.5 seconds after odor onset) for the CS+ and CS-, in late training (days 11+), for exp. mice (colored dots) and ctrl. mice (gray dots). Black lines indicate means. Stars indicate the significance level of paired or unpaired t-tests. Cue responses were significantly larger in the exp. group than the ctrl. group for the CS+ (mean %dF/F ± SEM, exp. mice [*n* = 5]: 0.66 ± 0.33, ctrl. mice [*n* = 3]: −0.67 ± 0.04, *P =* 0.023, unpaired t-test) but not the CS-(mean %dF/F ± SEM, exp. mice [*n* = 5]: −0.37 ± 0.23, ctrl. mice [*n* = 3]: −0.62 ± 0.06, *P =* 0.46, unpaired t-test) (*Property 1: Cue Response*, **Fig. 1b**). Omission responses were significantly more negative in the exp. group than the ctrl. group for the CS+ (mean %dF/F ± SEM, exp. mice [*n* = 5]: −3.23 ± 0.44, ctrl. mice [*n* = 3]: −1.23 ± 0.11, *P =* 0.015, unpaired t-test) but not the CS-(mean %dF/F ± SEM, exp. mice [*n* = 5]: −1.21 ± 0.12, ctrl. mice [*n* = 3]: −1.13 ± 0.08, *P =* 0.67, unpaired t-test) (*Property 2: Omission Dip*, **Fig. 1b**). Within the exp. mice, cue responses were larger to the CS+ than the CS-(mean %dF/F ± SEM, exp. mice [*n* = 5], CS+: 0.66 ± 0.33, CS-: −0.37 ± 0.23, *P =* 0.0019, paired t-test) and omission responses were more negative (mean %dF/F ± SEM, exp. mice [*n* = 5], CS+: −3.23 ± 0.44, CS-: −1.21 ± 0.12, *P =* 0.018, paired t-test). j) Average GRAB_DA3m_ responses to unpredicted opto-stimulation (0-1 seconds after opto-stimulation) or predicted opto-stimulation (0-1 seconds after opto-stimulation on stimulated CS+ trials). Note the y-axis scale difference compared to panel i. Predicted opto-stimulation responses were smaller than unpredicted opto-stimulation responses (mean %dF/F ± SEM, exp. mice [*n* = 5], unpred: 19.52 ± 1.80, pred: 17.13 ± 2.01, *P =* 0.0051, paired t-test) (*Property 3: Expectation-dependent reduction of outcome response*, **Fig. 1b**). k) Schematic (left) and example histology image (right) showing insertion of a four-shank Neuropixels 2.0 probe into VTA (see also **Extended Data Fig. 1e**). l) Schematic of antidromic optotagging. Red light delivery in lNAc generates dopamine release (green) as well as back-propagating action potentials which travel antidromically along dopamine axons to VTA dopamine neuron cell bodies, where they are detected by the Neuropixels probe, marking the recorded neuron as a dopamine neuron that projects to lNAc. m) Example antidromically optotagged dopamine neuron (see Methods for criteria defining optotagged neurons). n) The response of four example antidromically optotagged lNAc-projecting VTA dopamine neurons to CS+ (omission trials; red) and CS-(black) (see also **Extended Data Fig. 2c**). o) The average response of all optotagged neurons to CS+ (omission trials) and CS-(mean ± SEM, *n* = 43 neurons). Firing rate traces were smoothed and then z-scored using the mean and standard deviation of the firing rate trace in ITI periods (Methods). (see also **Extended Data Fig. 2d**). p) Difference in z-scored response between the CS+ (omission trials) and CS-(mean ± SEM, *n* = 43 neurons). Cue period was defined as 0-0.5 seconds after odor onset, and omission period was defined as 0-0.5 seconds after omission of opto-stimulation (1.5-2.0 seconds after odor onset); these are used for quantification in panel q. q) The average cue and omission response (difference in z-score between CS+, on omission trials, and CS-, in the time windows defined in panel p) for all optotagged neurons. Points indicate neurons, and colors indicate mice. The four example neurons in panel n are labeled. At the single neuron level, cue responses were significantly greater than zero (mean z-score difference ± SEM [CS+ − CS-] = 0.13 ± 0.04, *n* = 43 neurons, *P =* 0.0017, t-test), and omission responses were significantly less than zero (mean z-score difference ± SEM [CS+ − CS-] = - 0.19 ± 0.05, *n* = 43 neurons, *P =* 0.00054, t-test). These were marginally anti-correlated (Pearson’s *ρ* = −0.29, *P =* 0.06). In histograms, neurons with significant cue responses or omission dips are depicted with black bars, and non-significant neurons with open bars.

The TD algorithm relies on a core derivative-like computation, from which it derives its name: the comparison of value estimates at adjacent time steps (**Fig. 1a**). The intuition is simple: If value suddenly increases, then to the extent that this was predictable, previous states were undervalued, and vice versa. This empowers the algorithm to leverage its internal estimates to drive learning in the absence of reward feedback, using a temporally local computation to bridge events that are far apart in time.

A profound implication of this is that the time horizon of the TD learning algorithm—its temporal discount factor, *γ*—depends on the relative weighting of current and recent time in the local comparison of the value function at adjacent time steps. This suggests that microcircuit mechanisms underlying the derivative computation in the dopamine system may quantitively explain animals’ preference for immediate versus future rewards. Time discounting is a fundamental aspect of learning and decision making^12,13^ and may be a partly hardwired personality trait^14^, but its biological origins remain mysterious. The possibility of quantitatively linking dopamine circuits to time preference is tantalizing but unrealized.

While debates continue^15–18^, the canonical evidence supporting the TD RPE interpretation of dopamine is that dopamine neurons in trained animals (1) respond strongly to unexpected rewards, but less to expected rewards; (2) respond to reward-predicting sensory cues; and (3) become suppressed when expected rewards are omitted^1^ (**Fig. 1b**). In addition to these canonical TD RPE-like activity patterns, it was recently discovered that ramping dopamine activity, which can occur before reward delivery^19,20^, reflects the approximate derivative of a hidden function, and thus can also be interpreted as a TD error^5,21,22^ (**Fig. 1b**). Yet despite these and other pieces of correlational^2,4^ and behavioral^3,23,24^ evidence, the biological mechanisms that produce TD error-like activity in dopamine neurons, including the core derivative-like computation, remain unknown.

We hypothesized that TD RPE-like activity in dopamine neurons arises through bidirectional interactions between midbrain dopamine neurons and their primary output region, the striatum—a longstanding idea^25–28^ that remains unproven. Neural activity in the striatum correlates with value^29–32^ and influences dopamine signaling both directly and indirectly, via its two primary cell types, D1 and D2 medium spiny neurons (MSNs) (**Fig. 1c**). However, the idea that dopamine-striatum interactions generate TD RPE signals in dopamine neurons has been difficult to test experimentally, for two reasons: (1) tasks trained with natural rewards engage learning processes throughout the entire brain, not just the dopamine system, so signaling patterns in dopamine neurons in natural tasks cannot be assumed to arise specifically from dopamine-striatum interactions; and (2) dopamine neurons are themselves diverse, with distinct properties and functions depending in part on projection target^33–37^, so dopamine neurons may not all participate in value learning.

To unravel this complexity step-by-step, we started with an artificial opto-conditioning task with just two components: sensory stimuli (odors) and optogenetic stimulation of dopamine axons in the lateral nucleus accumbens (lNAc), a subregion of the striatum with concentrated TD RPE-like dopamine signaling^38,39^ and value-related neuronal activity^31,32^ in natural conditioning tasks. The results revealed that dopamine neurons are hardwired to automatically compute a temporal difference of input from D1-MSNs, resulting in patterns of TD error in dopamine signaling, and suggest that the balance of excitation and inhibition in this pathway sets the temporal discount factor of the TD algorithm.

## Results

### Opto-conditioning in lNAc generates TD error signals in dopamine activity

Mice were prepared for opto-conditioning by virally expressing red excitatory opsin ChrimsonR^40^ unilaterally in ventral tegmental area (VTA) dopamine neurons and green dopamine sensor GRAB_DA3m_^41^ in lNAc (**Fig. 1d**). An optic fiber was implanted targeting lNAc in the same hemisphere, through which dopamine axons could be stimulated while simultaneously recording dopamine release using photometry without optical crosstalk (**Extended Data Fig. 1a,b**) or movement artifacts (**Extended Data Fig. 1c**). To ensure physiological relevance, optogenetic stimulation was calibrated to a natural reward (8 *μ* droplet of water) (**Fig. 1e, Extended Data Fig. 1b**)^42^.

Mice underwent conditioning in which odors (CS+/CS-) preceded opto-stimulation (75% of CS+ trials) or no outcome (100% of CS-trials, 25% of CS+ trials) (**Fig. 1f, Extended Data Fig. 1d,e**). In contrast to a previous report^16^, lNAc dopamine in ChrimsonR mice (*n* = 5), but not control tdTomato mice (*n* = 3), gradually developed classic signatures of TD error (**Fig. 1g-j, Extended Data Fig. 1f-i**): a positive response to CS+ (“cue response”), preceding the time of predicted stimulation (**Fig. 1g-i**); a negative response aligned to omission of predicted stimulation (“omission dip”) (**Fig. 1g-i**), which reflected the timing of stimulation used during training (**Extended Data Fig. 1j-l**); and an expectation-dependent reduction in the opto-stimulation response (**Fig. 1j**).

To investigate these changes at the level of single dopamine neurons while preserving projection target specificity, we recorded spiking activity of antidromically optotagged lNAc-projecting VTA dopamine neurons in the same mice on a subset of sessions using Neuropixels probes (**Fig. 1k-q, Extended Data Fig. 2**). Projection target was identified by stimulating dopamine axons in lNAc while recording spikes at the cell body in VTA (**Fig. 1k-m**). As in the photometry data, cue responses and omission dips could be detected in the spiking activity of individual dopamine neurons (**Fig. 1k-q, Extended Data Fig. 2**). While statistically significant both within and across mice, the cue response and omission dip were small in both spiking and photometry signals (∼5% of response to uncued water delivery, **Fig. 1i,j, Extended Data Fig. 2**)^16^, likely reflecting the minimal nature of our calibrated, local, unilateral dopamine axon stimulation, and/or possibly a missing factor that would normally accompany a natural reward.

Opto-conditioning was sufficient to alter behavior: Mice exhibited greater anticipatory facial movements following the CS+ than the CS-(**Extended Data Fig. 3**). However, behavioral and neural changes were dissociable, as neural changes were observed even in mice that exhibited no detectable change in movement following odor onset.

### Opto-conditioning in lNAc generates a value-like signal in D1-but not D2-MSNs

To understand how opto-conditioning in lNAc generated TD error-like signals in dopamine neurons, we used Neuropixels probes to record extracellular spiking activity in the striatum, on a subset of late-training days in the same mice (**Fig. 2a**). We selected neurons with a positive response to either the CS+ or CS-(Methods) and plotted their activity on omission trials. In contrast to the biphasic response observed in dopamine neuron spiking (**Fig. 1o,p**) and dopamine release (**Fig. 1g,h**), striatal spiking was positively enhanced to the CS+ relative to the CS-throughout the trial (**Fig. 2b, Extended Data Fig. 4**), reminiscent of a “value”-like signal as opposed to the TD error-like patterns observed in dopamine activity^43^. This change was observed specifically within lNAc (the stimulation site) and not dorsal striatum (DS) (**Fig. 2c**), indicating that opto-conditioning was confined to the target region (lNAc). Selecting neurons inhibited by either odor did not reveal a clear enhancement of inhibition to the CS+; if anything, these neurons’ responses were also potentiated (**Extended Data Fig. 4e**), suggesting that the primary effect of opto-conditioning was to positively enhance odor-evoked spiking in a subset of neurons.

**Figure 2:**
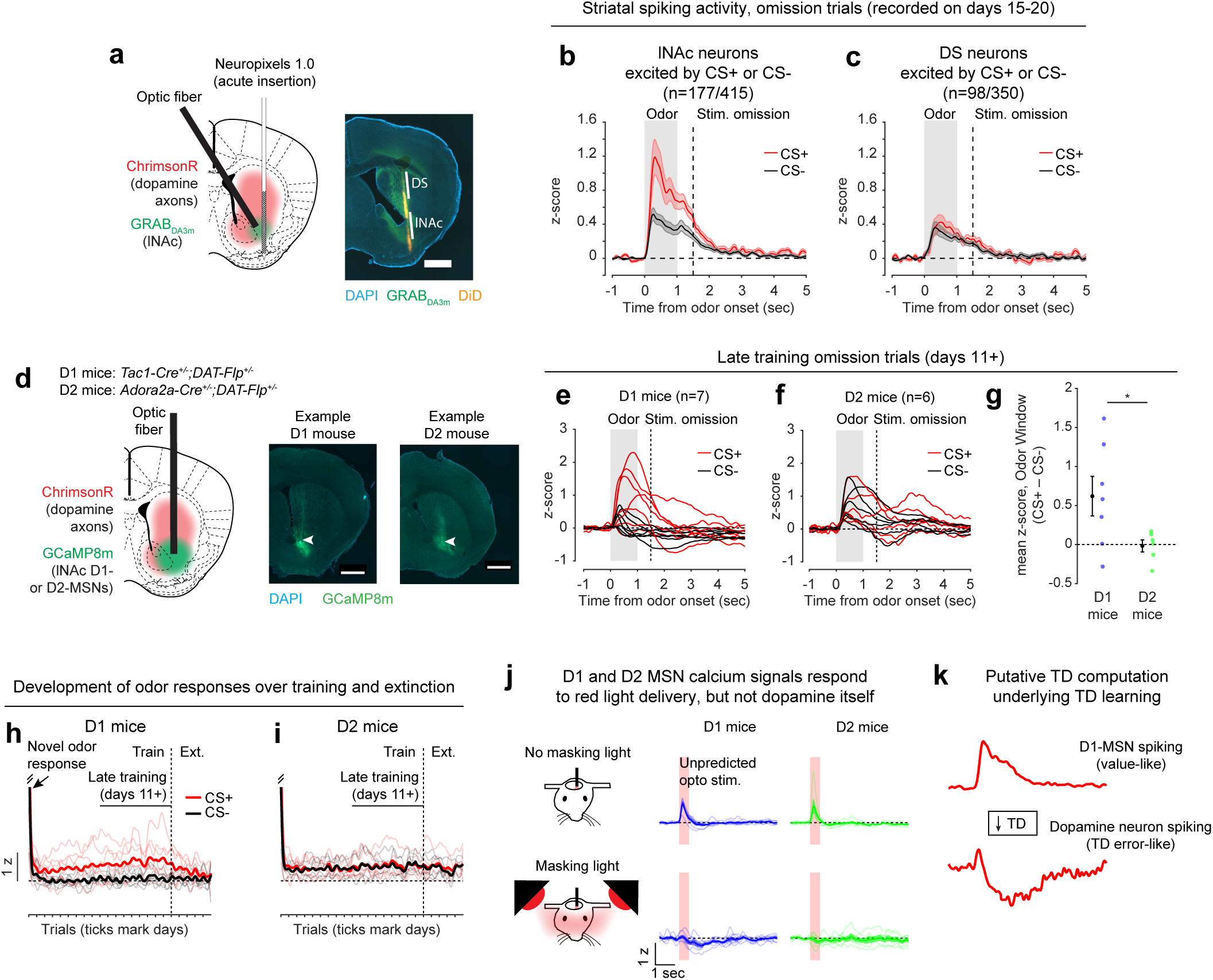
Opto-conditioning in lNAc generates a value-like signal in D1-but not D2-MSNs. a) Left: Schematic of Neuropixels 1.0 recording targeting lNAc during the opto-conditioning task. Right: Example histology image with lNAc (defined as 3.5-4.5 mm from brain surface) and dorsal striatum (DS; defined as 2-3 mm from brain surface) marked along the Neuropixels probe path. Scale bar = 1 mm. lNAc Neuropixels recordings were performed in the same mice as shown in **Figure 1**, on days 15-20 of training (**Extended Data Fig. 1e**). b) Mean z-scored striatal spiking response of lNAc odor-responsive neurons to the CS+ (omission trials; red) and CS-(black). Lines and shaded areas show mean ± SEM over neurons (*n* = 177 odor-responsive neurons). Odor-responsive neurons were defined as those having a significantly elevated firing rate relative to baseline in any 1-second bin from 0-5 seconds after odor onset, to either the CS+ or the CS-(177/415 neurons; Methods). Odor responses (defined as the average z-scored activity from 0-1 second after odor onset) of odor-responsive lNAc neurons were significantly greater for the CS+ than the CS-, at both the neuron and mouse level (mean z-score ± SEM, neurons: CS+ = 0.78 ± 0.13, CS-= 0.36 ± 0.05, *n* = 177 neurons, *P =* 6.2×10^-6^, sign-rank test; mice: CS+ = 0.75 ± 0.13, CS-= 0.29 ± 0.08, *n* = 5 mice, *P =* 0.032, t-test). c) Same as panel b, but for DS neurons. Unlike lNAc neurons, odor-responsive DS neurons did not differ in their response to the CS+ versus the CS-, at the neuron or mouse level (mean z-score ± SEM, neurons: CS+ = 0.30 ± 0.06, CS-= 0.25 ± 0.04, *n* = 98 neurons, *P =* 0.41, sign-rank test; mice: CS+ = 0.45 ± 0.24, CS-= 0.30 ± 0.10, *n* = 5 mice, *P =* 0.34, t-test). d) Left: Schematic of experimental approach to record from D1- and D2-MSNs during opto-conditioning. Using double transgenic mice, Flp-dependent ChrimsonR was expressed in VTA dopamine neurons and Cre-dependent GCaMP8m was expressed in lNAc D1-MSNs (*Tac1-Cre^+/-^;DAT-Flp^+/-^* mice, i.e. “D1 mice”, *n* = 7) or D2-MSNs (*Adora2a-Cre^+/-^;DAT-Flp^+/-^* mice, i.e. “D2 mice”, *n* = 6). An optic fiber was implanted targeting lNAc, through which dopamine axons could be stimulated while recording bulk calcium signals from D1- or D2-MSNs. Opto-stimulation was calibrated to water reward in separate *DAT-Flp^+/-^* mice (**Extended Data Fig. 5b**). Right: Example histology images showing fiber placement in lNAc (arrows) and GCaMP8m expression. Scale bars = 1 mm. e) Average z-scored bulk calcium response of D1-MSNs to CS+ (omission trials) and CS-in late training (days 11+) of the same opto-conditioning task as in **Fig. 1** (*n* = 7 D1 mice). For each mouse, there is one red line (CS+ response on omission trials) and one black line (CS-response). f) Same as panel e, but for D2 mice (*n* = 6). g) Average difference in z-score between CS+ (omission trials) and CS-for D1 mice (blue) and D2 mice (green) during odor delivery (0-1 seconds after odor onset). For D1 mice, this difference was greater than zero (marginally significant) (mean z-score ± SEM = 0.62 ± 0.25, *n* = 7, *P =* 0.0504, t-test), but for D2 mice it was not (mean z-score ± SEM = −0.02 ± 0.08, *n* = 6, *P =* 0.84, t-test), and the D1 odor response was significantly greater than the D2 odor response in a head-to-head comparison (*P =* 0.047, unpaired t-test). h) Development of odor responses (average z-score GCaMP signal 0-1 seconds after odor onset) to CS+ (red) and CS-(black) in D1 mice over training (“train”) and extinction (“ext.”). Single trial responses were averaged in the given time window, concatenated across days, and smoothed with a Gaussian filter (S.D. = 15 trials). Thin lines represent individual mice; thick lines show the mean over mice. i) Same as panel h, but for D2 mice. j) D1- and D2-MSN calcium signals respond to red light delivery, but not dopamine itself. Top row: z-scored GCaMP responses to unpredicted opto-stimulation (shaded red area) in D1 mice (left) and D2 mice (right). Ten unpredicted opto-stimulation trials were delivered at the end of the opto-conditioning task each day (**Extended Data Fig. 1e**). The average response to these trials is shown, averaged first within day then across days for each mouse (mean z-score ± SEM, 0-1 sec after light onset: D1 mice = 0.26 ± 0.06, *n* = 7, *P =* 0.0062, t-test vs zero; D2 mice = 0.20 ± 0.09, *n* = 6, *P =* 0.071, t-test vs zero). Each thin line shows an individual mouse; thick lines and shaded areas show means ± SEM over mice. Bottom row: Several days after the completion of opto-conditioning (including extinction sessions), a session was run in which unpredicted opto-stimulation was delivered while the recording rig was illuminated with two bright red lamps directed at the mouse’s eyes. The response to unpredicted opto-stimulation was eliminated in both D1 and D2 mice (mean z-score ± SEM, 0-1 sec after light onset: −0.12 ± 0.09, *n* = 7, *P =* 0.24, t-test vs zero; D2 mice = −0.09 ± 0.06, *n* = 6, *P =* 0.22, t-test vs zero; paired t-test vs no masking light: *P =* 0.0012, D1 mice; *P =* 0.0096, D2 mice), indicating that these responses were visual rather than a response to dopamine itself. k) Putative TD computation underlying TD learning. Opto-conditioning in lNAc potentiates lNAc D1-MSN spiking to CS+ relative to CS-, generating a value-like signal reflecting a prediction of upcoming dopamine stimulation but not dopamine stimulation itself. Monophasic lNAc D1-MSN spiking activity undergoes a temporal difference transformation to produce biphasic TD error-like patterns of VTA dopamine spiking activity (and lNAc dopamine release) in response to the CS+.

We next aimed to determine which striatal cell types contribute to the changes in striatal spiking generated by opto-conditioning. The two primary striatal cell types are D1- and D2-MSNs, expressing the D1 and D2 dopamine receptor, respectively. We used double transgenic mice to virally express a calcium indicator (GCaMP8m) in lNAc D1- or D2-MSNs along with ChrimsonR in VTA dopamine neurons (*n* = 7 D1 mice, *n* = 6 D2 mice; **Fig. 2d, Extended Data Fig. 5a,b**; Methods). These mice then underwent the same two-odor opto-conditioning paradigm as above. Consistent with asymmetric plasticity rules in D1- and D2-MSNs^44–46^, phasic dopamine stimulation selectively potentiated D1-MSN but not D2-MSN responses to preceding odor stimuli (**Fig. 2e-i, Extended Data Fig. 5c-f**). We conclude that the changes in lNAc spiking activity (**Fig. 2b**) primarily reflect changes in D1-MSNs.

A key feature of a value-like signal is that it reflects a prediction of upcoming reward but does not respond directly to reward itself. Consistent with this, D1-MSN odor responses gradually increased over training (**Fig. 2h**), but D1-MSNs did not respond directly to dopamine release (**Fig. 2h, Extended Data Fig. 4c**)^47^. We conclude that the D1-MSN response to the conditioned odor is “value-like” in that it reflects a gradually learned prediction of upcoming dopamine stimulation but does not respond to dopamine itself.

Together, these activity patterns are suggestive of a TD learning loop: Dopamine release gradually potentiates the odor-evoked response in D1-MSNs (“value”); in turn, D1-MSNs stimulate lNAc-projecting VTA dopamine neurons via a temporal difference transformation, generating a positive dopamine response at onset and a negative omission dip at offset of their activity, closely recapitulating TD RPE (**Fig. 2k**).

### Activation of lNAc D1-MSNs drives lNAc dopamine release via a TD transformation

To directly test this idea, we asked whether activation of D1-MSNs alone could generate the TD error-like patterns of dopamine activity observed in our opto-conditioning task, and more generally, whether D1-MSNs drive dopamine release according to a temporal difference transformation—the core computation in TD learning (red box, **Fig. 1a**). We prepared mice for simultaneous opto-stimulation of lNAc D1-MSNs with photometric recording of lNAc dopamine release and Neuropixels recordings of lNAc D1-MSNs (**Fig. 3a**) and designed 7 opto-stimulation patterns that generate both qualitative and quantitative predictions for a TD error-like response (**Fig. 3b**): (1) A 1 second pulse train at 20 Hz, (2) a 2 second pulse train at 20 Hz, (3) an upward ramp from 0 to 40 Hz, (4) a downward ramp from 40 to 0 Hz, and (5-7) 3 second pulse trains at 5, 10, and 20 Hz (all light pulses 1 ms at 40 mW/mm^2^).

**Figure 3:**
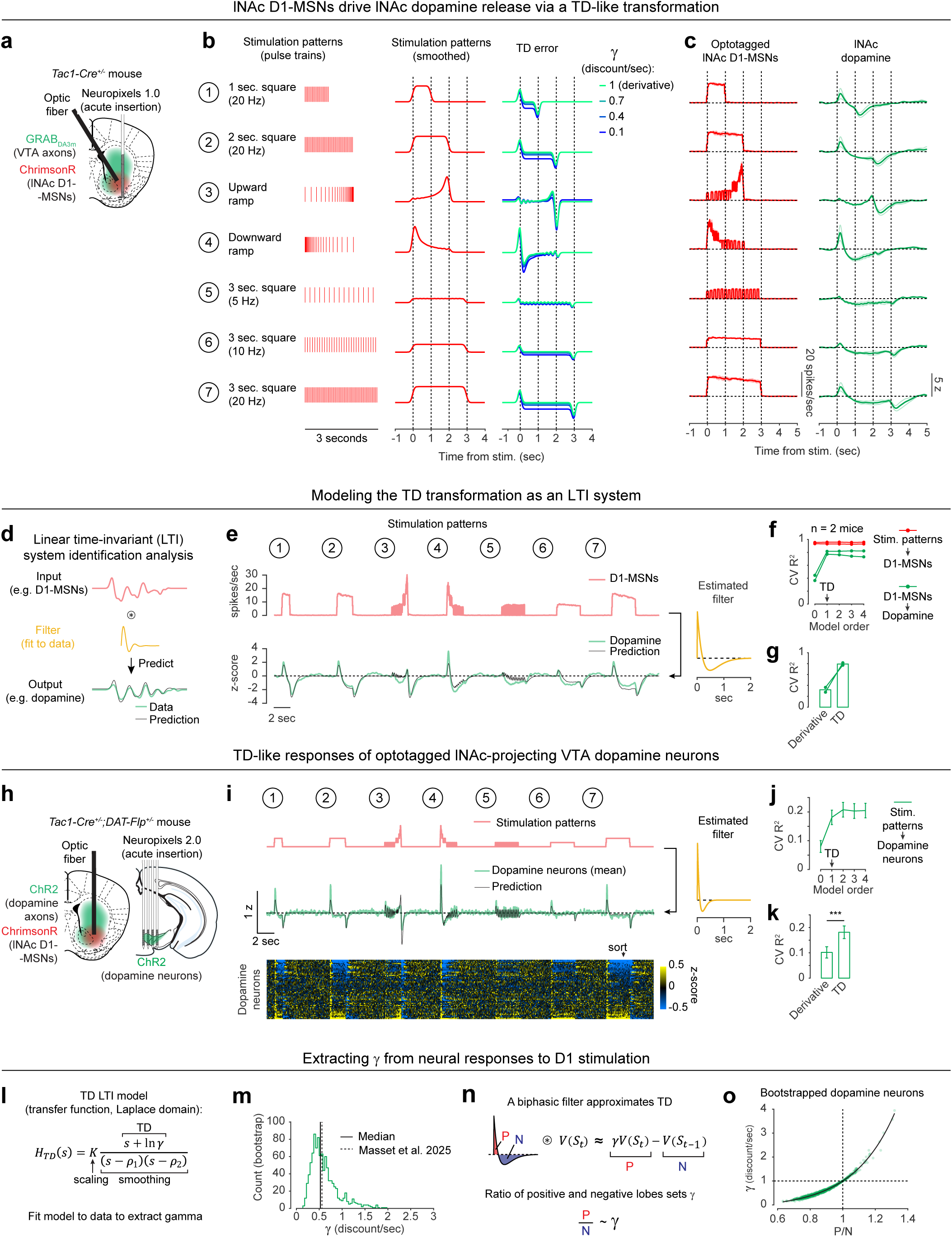
Activation of lNAc D1-MSNs drives lNAc dopamine neurons via a temporal difference transformation, suggesting a circuit-level origin of temporal discounting. a) Experimental approach. GRAB_DA3m_ was expressed pan-neuronally in the VTA and ChrimsonR was expressed in D1-MSNs in lNAc using *Tac1-Cre^+/-^*mice. An optic fiber was implanted at an angle targeting lNAc allowing for acute insertion of a Neuropixels probe as in **Fig. 2a**. This enabled simultaneous stimulation of lNAc D1-MSNs, electrophysiological recording of optotagged lNAc D1-MSNs, and photometric recording of lNAc dopamine release from VTA axons at the same site, such that the transfer function from D1-MSN activity to lNAc dopamine release could be directly quantified. b) *Left:* The 7 opto-stimulation patterns used in the experiment (trains of 1 ms red light pulses at 40 mW/mm^2^). *Middle:* Smoothed pulse trains. *Right:* The TD equation applied to the smoothed pulse trains, with varying *γ*. When *γ* = 1, TD becomes a pure derivative. For *γ* < 1, TD deviates from a derivative, most notably in its negative steady-state response (e.g. Patterns 1, 2, 7). c) The responses of optotagged lNAc D1-MSNs (left) and lNAc dopamine (right) to the 7 stimulation patterns (*n* = 2 mice). Individual D1-MSNs were averaged within mice (*n* = 6, 10 D1-MSNs in mouse 1, 2 respectively). Faint lines show individual mice, solid lines show the average over mice. Whereas D1-MSN activity closely followed the stimulation patterns themselves, dopamine release qualitatively resembled TD error of the stimulation patterns, including a negative steady-state response, which distinguishes TD from a pure derivative. d) Schematic of linear time-invariant (LTI) systems identification analysis, which models an input-output system as a linear filter (also called the impulse response) convolved with the input to produce the output (Methods). e) LTI model fit to data from panel c. *Top*: Average lNAc D1-MSN responses to the 7 stimulation patterns (red; averaged over *n* = 2 mice). *Bottom*: The average response of lNAc dopamine to the 7 stimulation patterns (green; averaged over *n* = 2 mice) and the LTI model fit (black) predicting lNAc dopamine (output) from opto-triggered D1-MSN activity (input) (cross-validated [CV] R^2^ = 0.81). *Right:* Estimated filter of LTI model (Methods). f) Cross-validated R^2^ for LTI models of varying order (number of zeros; Methods), predicting D1-MSN activity from stimulation patterns (red) or dopamine activity from D1-MSN activity (green). Individual lines show individual mice (*n* = 2). The TD equation is order 1 (Methods; panel l). Consistent with a TD transformation between D1-MSNs and dopamine, predicting dopamine from D1-MSNs required a minimum model order of 1, and higher model orders did not improve performance. In contrast, an order 0 model was sufficient to predict D1-MSN activity from stimulation patterns. g) A derivative LTI model performed worse than a TD LTI model at predicting dopamine from D1-MSNs. The derivative model is equivalent to the TD LTI model with *γ* fixed at 1 (Methods). h) Experimental design to record from optotagged lNAc-projecting VTA dopamine neurons with Neuropixels probes while optogenetically stimulating lNAc D1-MSNs. i) *Top*: The 7 stimulation patterns, concatenated. *Middle:* Average dopamine neuron response (z-score) to 7 stimulation patterns (green) and the LTI model fit (black) predicting dopamine from stimulation patterns. Note the qualitative similarity to the bottom of panel e (CV R^2^ = 0.69). *Bottom:* Heatmap of activity of single optotagged lNAc-projecting VTA dopamine neurons, sorted by the response to pattern 7 (*n* = 55 neurons). *Right:* Estimated filter of LTI model fit to average dopamine neuron spiking activity. Note the temporally sharper shape compared to panel e, befitting the higher temporal resolution of electrophysiological recording. j) Same as panel f, but for models predicting single optotagged dopamine neuron activity from stimulation patterns. Lines and error bars indicate mean ± SEM (*n* = 55 neurons). k) Same as panel g, but for models predicting single optotagged dopamine neuron activity from stimulation patterns. *** *P <* 0.001, paired t-test (*n* = 55 neurons). l) Schematic describing how the temporal discount factor *γ* can be extracted from dopamine responses to D1-MSN stimulation using an order-1 LTI model (Methods). m) Histogram of estimated *γ* from bootstrapped optotagged dopamine neuron population activity (1000 bootstraps; Methods). The median bootstrapped estimate (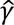 = 0.53) closely matched the estimate of *γ* from a previously published paper using a cued delay task (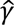 = 0.56, Masset et al. 2025^48^; c.f. Sousa et al. 2025^49^ and Kobayashi & Schultz 2008^50^). n) Schematic demonstrating how convolution with a biphasic filter approximately implements a TD transformation, and the intuition for why the ratio of the filter’s positive and negative lobes (P and N respectively) should set the temporal discount factor, *γ*. o) The ratio of the positive and negative lobes of the fit impulse response (P/N) versus the parameter *γ* of the LTI fit estimated as in panel l. Green points show individual bootstrapped estimates (Methods) and the black line shows an exponential fit.

The simulated TD error of these 7 patterns is shown in the right of **Fig. 3b**, with the temporal discount factor *γ* ranging from 1 (green; a pure derivative) to 0.1 (blue) in units of discount/sec. As has been previously noted^5,21,22^, TD error is similar (but not identical) to a time derivative. Indeed, in the schematics in **Fig. 3b**, the TD error shows several qualitative derivative-like features that are shared between TD in general (*γ* ≤ 1) and the strict derivative (*γ* = 1): (1) Transient positive/negative responses to the onset/offset of stimulation, which scale with the intensity of stimulation (patterns 5-7), (2) A prominent negative response to the offset of pattern 3, with a smaller positive response during the ramp-up phase, and (3) A prominent positive response to the onset of pattern 4, followed by a prolonged negative response as stimulation ramps down.

Remarkably, actual dopamine responses to D1-MSN stimulation displayed all of these derivative-like features (**Fig. 3c**, right). In contrast, D1-MSNs responded faithfully to the stimulation patterns themselves (**Fig. 3c**, left; **Extended Data Fig. 6a**). Notably, even though striatal MSNs are inhibitory neurons and directly project to VTA, the initial dopamine response was excitatory, especially for stimulation patterns with a rapid onset. This counterintuitive response pattern supports the existence of a derivative-like transformation between lNAc D1-MSNs and lNAc dopamine release, exactly as needed to implement TD learning (**Fig. 1a**).

While TD error is derivative-like, it deviates from a derivative when *γ* < 1, most prominently in the steady-state response to sustained stimulation: The steady-state response is zero for *γ* = 1 (derivative) but negative for *γ* < 1. Dopamine responses better matched TD, with negative steady state responses that scaled with stimulus intensity (**Fig. 3c**). Computationally, this key distinction gives the TD algorithm its preference for current over future rewards. These data raise the possibility that the temporal discount parameter *γ* could be set by a neural circuit mechanism between D1-MSNs and dopamine neurons—a possibility we explore in greater depth below.

### Capturing the TD transformation as convolution with a biphasic linear filter

In theory, a TD transformation can be viewed as convolution with a *biphasic filter,* which applies positive weights to recent input, negative weights to preceding input, and sums these to generate the output (**Fig. 3n**)^25^. To describe our results quantitatively, we therefore used linear time-invariant (LTI) systems identification analysis, a standard approach in engineering^51^, to estimate the linear filter that best captures the transformation from an input (e.g. D1-MSN spiking) to an output (e.g. lNAc dopamine release) (**Fig. 3d**) (Methods).

Whereas the best-fit filter from stimulus to D1-MSN spiking resembled a delta function, scaling the input to produce the output (**Extended Data Fig. 6d**), the best-fit filter from D1-MSN spiking to dopamine release was biphasic, with a rapid positive component followed by a delayed negative component, thus implementing an approximate temporal difference (**Fig. 3e**; order 1 LTI model; Methods). The LTI models accurately predicted held-out data for both transformations (cross-validated R^2^: stimulation patterns → D1-MSNs = 0.95, D1-MSNs → dopamine release = 0.81). These results argue that our data can be adequately understood as the result of a linear system implementing a TD transformation between D1-MSNs and dopamine release.

To examine how well the dopamine response matches TD as opposed to more complex models, we fit LTI models of varying order, using a standard model formulation which assumes a particular functional form of the filter (Methods)^51^. Briefly, in this formulation the *n*th order LTI model is composed of *n* TD transformations, applied sequentially, followed by smoothing and scaling. Each TD transformation is given by convolution with the “TD filter,” ℎ_*TD*_(*t*):

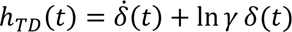

Where *δ*(*t*) and 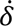(*t*) are the Dirac delta function and its derivative, respectively. Convolving an input *V*(*t*) with this filter generates an output 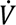(*t*) + ln *γ V*(*t*), which is exactly the continuous form of the TD equation^21^. The TD LTI model is order 1: It applies exactly 1 TD transformation, followed by smoothing and scaling. In contrast, the zeroth order model applies smoothing and scaling but lacks a TD transformation, whereas higher order models apply multiple TD transformations before smoothing and scaling, enabling higher order derivatives of the input to influence the output.

To capture the transformation from D1-MSNs to dopamine release, an order 1 LTI model was required, but higher order models did not improve performance (green lines, **Fig. 3f**). In contrast, a zeroth order LTI model was sufficient to capture the transformation from the stimulation patterns to D1-MSNs (red lines, **Fig. 3f**). This tightens the link between dopamine responses and the TD equation: simpler models are insufficient, and more complex models are not needed.

Next, we used the LTI framework to demonstrate that dopamine responses are better fit by the TD equation than by a pure derivative. We refit the order 1 LTI model while fixing *γ* = 1, thus implementing a pure derivative, as ln(1) = 0. This model performed substantially worse on held-out data than the order 1 LTI model in which *γ* was free to vary (**Fig. 3g**).

The pattern of dopamine results was replicated in a larger cohort without D1-MSN recordings, receiving only patterns 1-4 (*n* = 13 mice; **Extended Data Fig. 6e,h,k**), and no dopamine responses were observed in mice with tdTomato instead of ChrimsonR in D1-MSNs (*n* = 6 mice; **Extended Data Fig. 6f,i,l**), ruling out optical artifacts or visual responses. Mice with GFP instead of GRAB_DA3m_ in VTA axons showed slow biphasic fluctuations in GFP fluorescence following opto-stimulation, potentially reflecting a hemodynamic response, but GFP signals were small compared to GRAB_DA3m_ (*n* = 3 mice; **Extended Data Fig. 6g,j,m**).

### TD-like response in optotagged lNAc-projecting VTA dopamine neurons

To gain insight into the TD computation at the level of single neurons in the VTA and better estimate the rapid dynamics of the dopamine response, we recorded spiking signals of antidromically optotagged lNAc-projecting VTA dopamine neurons while stimulating lNAc D1-MSNs with the same 7 stimulation patterns (**Fig. 3h**). The same TD-like patterns as in **Fig. 3c** were observed in dopamine spiking activity (**Fig. 3i**), well-captured by a biphasic filter with a sharper time course (cross-validated R^2^ = 0.69; **Fig. 3i**, right). Like photometry signals, single dopamine neurons required an order 1 LTI model, but not higher (**Fig. 3j**), and were better fit by TD than a pure derivative model (**Fig. 3k**). The latency of the onset of the excitatory response to pattern 4 was less than 50 ms, and the response peaked between 50 and 100 ms (**Extended Data Fig. 6c**). Importantly, these data localize the TD-like transformation downstream of D1-MSN spiking but upstream of VTA DA neuron spiking, and constrain the time course of activation, narrowing down potential mechanisms.

### Extracting the temporal discount parameter *γ* from dopamine neuron responses to D1-MSN stimulation

The LTI modeling framework allows us to extract the temporal discount parameter *γ* directly from dopamine responses to D1 stimulation (**Fig. 3l**). We derived an exact mathematical relationship between *γ* and a parameter of the LTI model, namely *γ* = *e*^−*σ*^, where *σ* is the zero of the order 1 LTI model fit (Methods). Bootstrapped estimates of *γ* from optotagged dopamine neurons closely matched a previous estimate, measured in mice performing a cued delay task^48–50^ (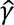 = 0.53 discount/sec in our data, 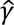= 0.56 discount/sec in Masset et al.^48^; **Fig. 3m**; note, however, that in Sousa et al.^49^ 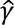≈ 0.9). *γ* in the LTI model is linked to the ratio of the positive and negative lobes of the impulse response (P/N) via an exponential relationship (**Fig. 3n,o**). This points to the relative magnitude and timing of the net-excitatory and net-inhibitory components of the dopamine response to D1-MSN activation as a key determinant of *γ*, grounding a higher order cognitive parameter (the time horizon of reward preference) in a concrete neural circuit mechanism.

### lNAc contributes a value-like signal to VTA dopamine

Because these experiments relied exclusively on artificial paradigms, we next asked whether lNAc contributes to dopamine TD RPE signaling in tasks with natural rewards. GCaMP6f was expressed transgenically in dopamine neurons by crossing *DAT-Cre*^+/-^ and *Ai148*^+/+^ mice. We then prepared mice for injection of muscimol (a GABA_A_ receptor agonist) or saline into the lNAc with photometry recording in VTA (**Fig. 4a**). We chose to record somatic rather than axonal calcium signals because muscimol may directly inhibit dopamine axons, which express GABA receptors^52^.

**Figure 4:**
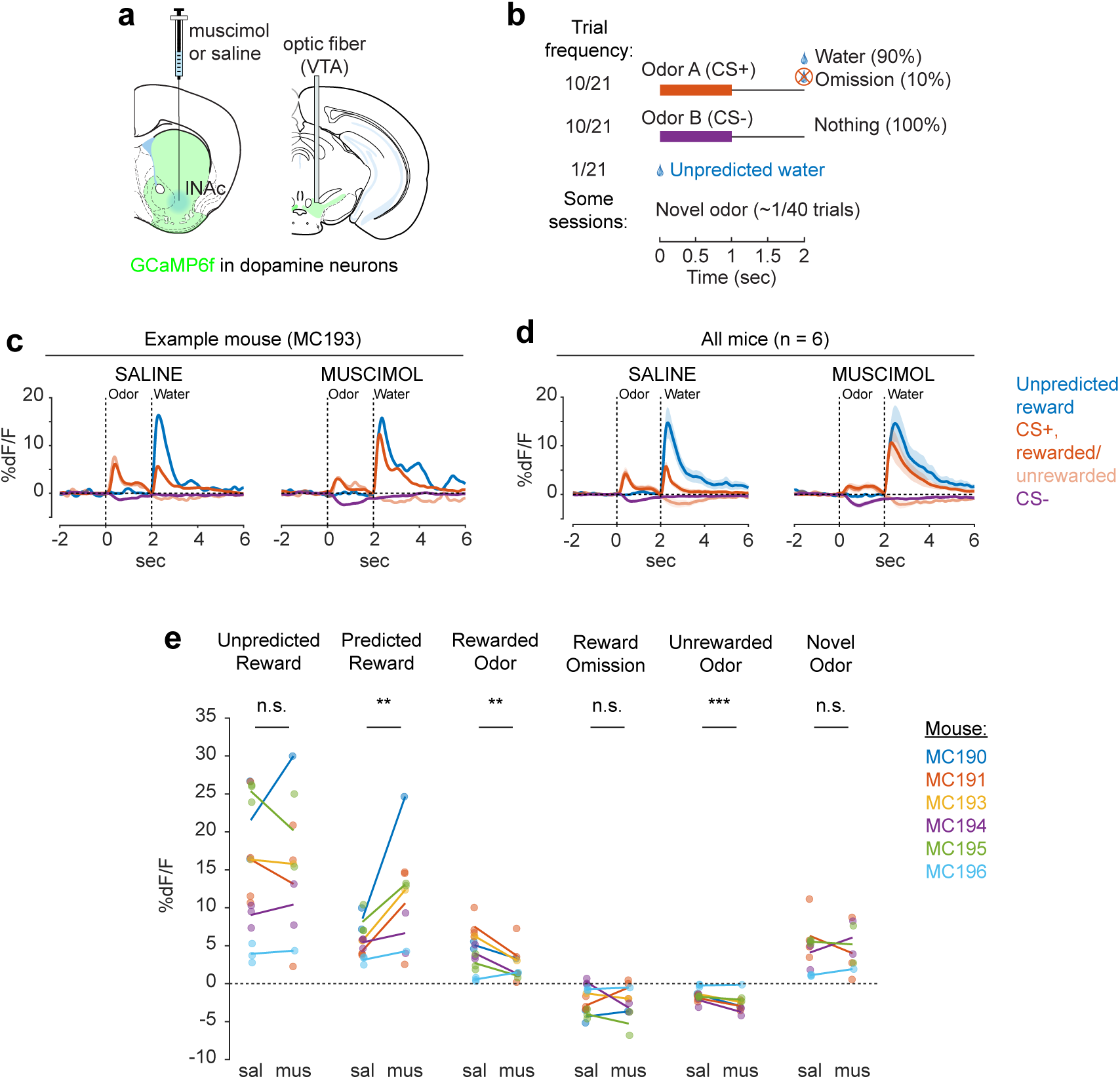
lNAc contributes a value-like signal to VTA dopamine neuron activity. a) Mouse preparation. GCaMP6f was expressed transgenically in dopamine neurons using *DAT-Cre^+/-^;Ai148^+/-^*mice (*n* = 6). An optic fiber was implanted targeting the VTA. After training in a classical conditioning task (panel b), muscimol or saline was injected into the lNAc on alternating days. b) Classical conditioning task design. c) Average VTA dopamine neuron calcium response to task events during example saline (left) and muscimol (right) sessions from the same mouse. Under muscimol, the response to the CS+ decreased and the response to the predicted reward increased, whereas the response to unpredicted reward remained relatively unchanged, consistent with lNAc signaling the reward expectation (value) associated with the CS+, and VTA dopamine neurons responding as the TD error of this value. Note that the response to the CS-also decreased, which may reflect generalization of value across odor stimuli within lNAc (e.g. the response to the CS-was positive in **Fig. 2b** and for some mice in **Fig. 2e**). d) Same as panel c, but averaged over mice (*n* = 6). Lines with shaded areas indicate mean ± standard error over mice for each trial type. e) Quantification of the traces in panel d. Points indicate the maximum (unpredicted reward, predicted reward, rewarded cue, novel odor) or minimum (reward omission, unrewarded cue) of the average %dF/F trace within a 1-second window following the given task event within individual sessions. Lines connect the average saline and average muscimol session for each mouse. Colors indicate mice. Stars indicate the significance of the injection type coefficient (saline vs. muscimol) in a linear mixed effect model with a random effect per mouse. Muscimol increased predicted reward responses (estimated muscimol coefficient [%dF/F] ± SEM = 5.47 ± 1.45, *n* = 26 sessions, *P =* 0.0013); decreased CS+ responses (−2.30 ± 0.67, *n* = 26 sessions, *P =* 0.0029); and decreased CS-responses (−0.93 ± 0.22, *n* = 26 sessions, *P =* 0.00046), but did not significantly affect unpredicted reward responses (−0.73 ± 2.23, *n* = 26 sessions, *P =* 0.75), reward omission responses (−0.21 ± 0.58, *n* = 26 sessions, *P =* 0.73), or novel odor responses (−0.28 ± 1.21, *n* = 21 sessions, *P =* 0.82). *** *P <* 0.001, ** *P <* 0.01, * *P <* 0.05, n.s. Not Significant.

Mice were water-deprived and trained in a classical conditioning task in which one odor (CS+) was associated with delivery of a natural reward (4 *μ* water; delivered on 90% of CS+ trials) and another odor (CS-) was associated with nothing (**Fig. 4b**). In addition, unpredicted rewards were delivered without a preceding odor on ∼5% of trials. After several days of training, we injected muscimol or saline into lNAc on alternating days while recording dopamine neuron calcium signals in the VTA. Muscimol dramatically increased calcium responses to predicted rewards, but not unpredicted rewards, and reduced responses to the CS+ (**Fig. 4c-e, Extended Data Fig. 7**). This pattern is consistent with a suppression of the reward expectation (value) associated with the CS+: the reward expectation of the CS+ decreases, so the cue response is suppressed, and at the same time the predicted (but not unpredicted) reward becomes more surprising, and so the reward response increases.

We tested for the specificity of this effect to odor value by also delivering novel odors in a subset of mice (4/6 mice). In contrast to the reduction in the dopamine response to the CS+, the response to novel odors did not change (**Fig. 4e**), arguing for a specific effect on odor value rather than novelty.

Because dopamine activity can reflect movements^53–55^, we analyzed face videos to check whether muscimol changed odor- and reward-triggered movements (**Extended Data Fig. 8**). Under muscimol, mice discriminated CS+ and CS-odors, consumed rewards, and detected novel odors (**Extended Data Fig. 8b**), suggesting that muscimol did not eliminate animals’ ability or motivation to perform the task. We did, however, detect changes in total mouth movement during the CS+ and after rewards and odor cues (**Extended Data Fig. 8b**), but muscimol similarly increased mouth movements to both predicted and unpredicted rewards, whereas VTA dopamine neuron calcium dramatically increased to predicted but not unpredicted rewards (**Fig. 4e**), indicating that movements cannot explain the main pattern of our results.

Together, these results demonstrate that lNAc contains a reward expectation (value)-like signal that contributes to TD RPE dopamine activity during classical conditioning with natural rewards, consistent with the TD error-like response of dopamine neurons to opto-stimulation of lNAc D1-MSNs.

### The temporal difference computation is a widespread feature of striatal-dopamine circuitry

Dopamine neurons have heterogenous functions depending in part on their projection target^33,39,53,56–58^, and may not all signal TD errors or participate in value learning. To investigate whether the TD computation identified here generalizes to striatal dopamine systems beyond lNAc, we stimulated D1-MSNs in lNAc, dorsomedial striatum (DMS), and dorsolateral striatum (DLS) while recording dopamine release in each site (**Fig. 5a,b, Extended Data Fig. 9**). To our surprise, the estimated impulse response for each stimulation site was biphasic (**Fig. 5c**), indicating that the temporal difference transformation is a ubiquitous feature of the circuitry connecting striatal D1-MSNs to midbrain dopamine neurons. However, the parameters of the impulse response, including the estimated temporal discount factor *γ*, varied by striatal subregion, with the estimated *γ* being closer to 1 for DMS and DLS than lNAc (though not significantly so; **Fig. 5d,e**). This points to a unified computational account of phasic dopamine’s role in learning as a prediction error machine, with parameters and input/output spaces that vary by striatal subregion^59–61^.

**Figure 5:**
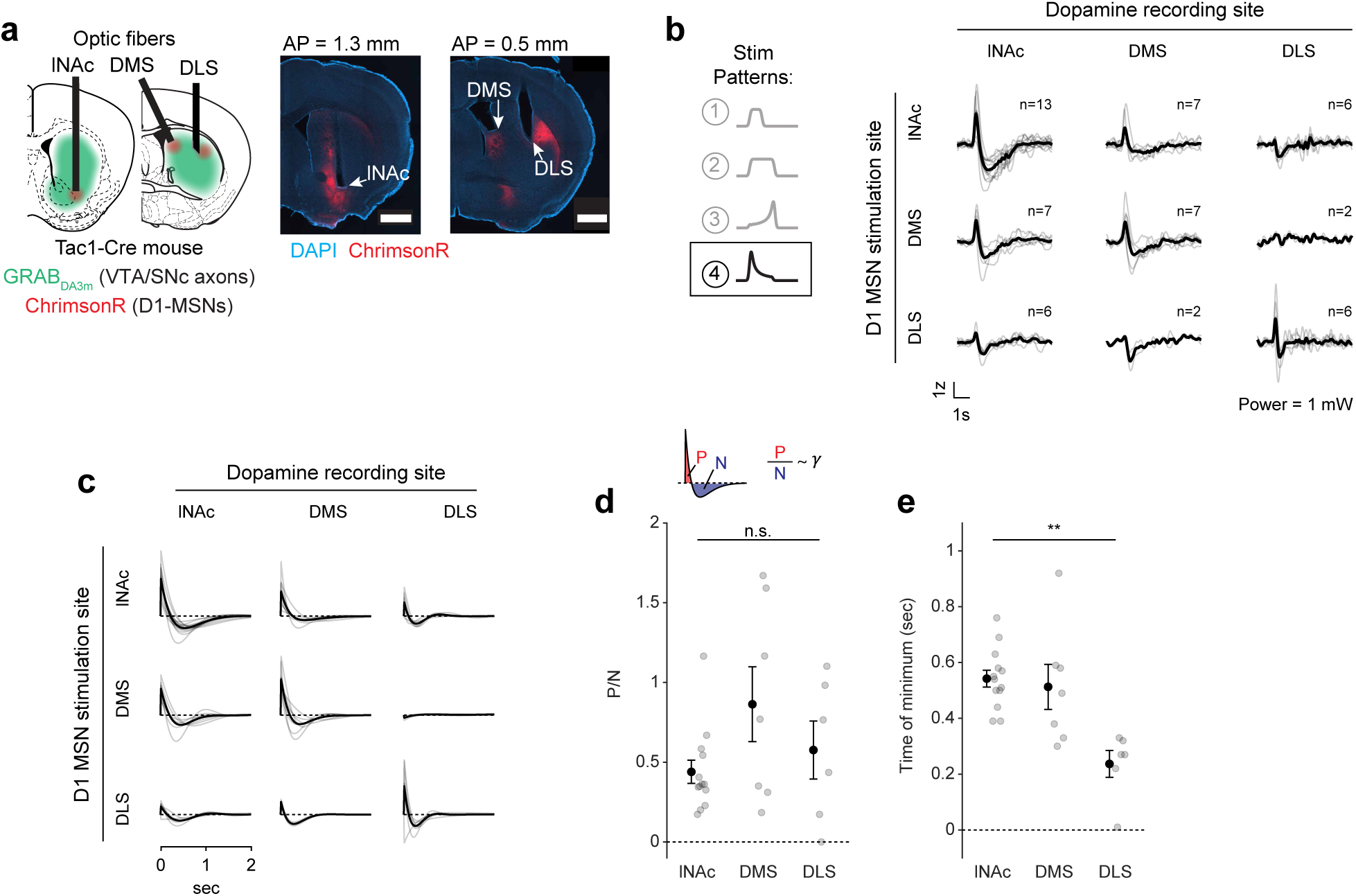
The temporal difference computation is a widespread feature of striatal-dopamine circuitry. a) Experimental design (left) and example histology images (right). GRAB_DA3m_ was expressed in VTA and SNc and ChrimsonR was expressed in D1-MSNs in lNAc, dorsomedial striatum (DMS), and/or dorsolateral striatum (DLS). Optic fibers were implanted targeting lNAc, DMS, and/or DLS. A mirrored fiber was used in DLS to restrict light delivery to DLS. Scale bar = 1 mm. b) Z-scored striatal dopamine responses to one stimulation pattern (Pattern 4, sharp onset followed by gradual downward ramp) (for responses to all patterns see **Extended Data Fig. 9**). The 3 x 3 matrix displays responses by dopamine recording site (columns) and D1-MSN stimulation site (rows). Each entry in the matrix contains a different number of mice (*n* = 13 mice total). Two mice had fibers in all three regions; the others had fibers in one or two of the three. All mice had fibers in lNAc. Gray lines = individual mice, black lines = mean over mice. Note the biphasic responses along the diagonal of the matrix, indicating a temporal difference computation for all three stimulation sites. c) Estimated impulse responses from LTI model predicting dopamine responses from stimulation patterns, as in **Fig. 3**, for all nine pairs of stimulation and recording sites. Gray lines = individual mice, black lines = mean over mice. Y-axis units are arbitrary. d) Ratio of the positive and negative areas of the impulse response (P/N, a proxy for *γ*; **Fig. 3n,o**) for lNAc, DMS, and DLS (when stimulating D1-MSNs in the same site; the diagonal of panel c). While estimated discount factors were closer to 1 for DMS and DLS compared to lNAc, this was not statistically significant (mean P/N ± SEM: lNAc = 0.50 ± 0.07, *n* = 13 mice; DMS = 0.94 ± 0.22, *n* = 7 mice; DLS = 0.65 ± 0.18, *n* = 6 mice; *P =* 0.23, linear mixed effects model). Gray points = individual mice, black points and error bars = means ± SEM over mice. e) Minimum time point of estimated impulse response for lNAc, DMS, and DLS (when stimulating D1-MSNs in the same site). Minimum time points were closer to zero for dorsal striatal areas, indicating faster dynamics^62,63^ (mean *t_min_* ± SEM [sec]: lNAc = 0.52 ± 0.03, *n* = 13 mice; DMS = 0.49 ± 0.08, *n* = 7 mice; DLS = 0.24 ± 0.05, *n* = 6 mice; *P =* 0.01, linear mixed effects model). Gray points = individual mice, black points and error bars = means ± SEM over mice. * *P <* 0.05, n.s. Not Significant.

## Discussion

Collectively, our experiments reveal how key TD learning operations are accomplished by the circuitry connecting lNAc-projecting dopamine neurons and lNAc D1-MSNs: phasic lNAc dopamine release selectively potentiates lNAc D1-MSN responses to preceding sensory stimuli, and D1-MSNs drive VTA dopamine neurons spiking and lNAc dopamine release via a TD transformation, generating TD error-like activity (a “minimal TD learning loop”; **Extended Data Fig. 10a**). As a counterpoint to recent arguments over whether TD RL is a good description of dopamine’s role in learning at the behavioral level^15–18^, these results demonstrate that TD error calculations are hard-wired into the dopamine system at the microcircuit level.

While the response of dopamine neurons to D1-MSN stimulation is derivative-like, it is not a strict derivative, but instead more closely matches the temporal difference equation *γV*(*S*_*t*_) − *V*(*S*_*t*−1_), with *γ* < 1. This is revealed most clearly by the response to long pulse trains (patterns 5-7, **Fig. 3c**), to which the dopamine response plateaus at a negative value that becomes more negative as stimulation frequency increases. In the LTI framework, this corresponds to a biphasic impulse response with total negative area (N) greater than total positive area (P). The ratio of these areas (P/N) determines the temporal discount parameter *γ* of the TD algorithm (**Fig. 3n,o**).

A natural extension of these findings is that the same neural circuit mechanism underlies the cognitive phenomenon of temporal discounting—the tendency for animals and humans to prefer immediate over future rewards. The time horizon of this preference—set by *γ*—is a fundamental aspect of learning and decision making with implications for human health and economic choices^13,64–67^. For instance, discount rates for money correlate with individuals’ choices to engage in unhealthy behaviors, such as smoking^66^ and excessive alcohol consumption^68^. The biological basis of time discounting is not known. Our findings suggest a compelling mechanism: *γ* is set at the level of individual dopamine neurons by the balance of the net-excitatory and net-inhibitory components of the circuit connecting D1-MSNs to VTA. This could explain why *γ* is a conserved property of single dopamine neurons across tasks^48^ and potentially explain the diversity of temporal discounting factors across striatal subregions^62,69^ and across individual animals in healthy and pathological conditions^66^. Notably, at the behavioral level temporal discounting is not exponential, as would be the case for a TD algorithm with a single discount factor, but hyperbolic, with a special status given to the current moment in time^12,50,70,71^ (but see^72^). A reservoir of dopamine neurons with private *γ*, each set individually by the balance the of positive and negative lobes of its impulse response to D1-MSN stimulation (**Extended Data Fig. 10b**), could combine to produce hyperbolic discounting on aggregate^48–50,73^. Further experiments linking the shape of the impulse response described here to neurophysiological and behavioral data in time discounting tasks would be needed to test this hypothesis (**Extended Data Fig. 10c**).

The TD transformation was a widespread feature of circuitry connecting D1-MSNs to dopamine release across both lNAc and dorsal striatum, revealing a conserved learning motif across striatal subsystems, despite their distinct roles in behavior^33,61^. This suggests that a single conserved TD-learning mechanism may have been repurposed by evolution to learn predictive maps across different input spaces^61,74^ and timescales^62,69^. Consistent with this, while the biphasic shape of the learned filter was conserved across the striatal areas we tested, the time resolution differed (time to minimum; **Fig. 5e**). The region-specific filters are likely shaped by a combination of both dopamine reuptake dynamics, which vary systematically across the striatum^63^, and circuit dynamics between striatum and dopamine neurons. Experiments involving projection-specific electrophysiological recording of the type developed here could distinguish between these alternatives.

Our experiments did not exhaustively cover striatal subregions, leaving several intriguing targets for future work, notably the medial nucleus accumbens, where dopamine release has value-like properties^39^, and the tail of the striatum, where dopamine may contribute to learning to predict threatening or salient outcomes^35,60,75^ or act as a value-neutral teaching signal^76^. If the hardwired TD computation extends to these regions, this would suggest dopaminergic learning follows an analogous algorithm despite notably different dopamine activity patterns. Outside striatum, the amygdala receives strong dopamine inputs from VTA, but appears to play a qualitatively distinct role in reinforcement learning compared to nucleus accumbens, with amygdala being more important for the initial acquisition of a behavioral response to motivationally salient cues (both rewarding and aversive)^77^, and nucleus accumbens subsequently refining the quantitative value estimate associated with these cues^78^. Distinct computations between amygdala/striatal neurons and dopamine neurons could help to explain this difference.

What are the possible biological mechanisms by which the temporal difference is computed? By recording lNAc D1-MSN optotagged neurons, antidromically optotagged lNAc-projecting VTA dopamine neurons, and lNAc dopamine photometry signals in the same paradigm, we show that the calculation occurs between the spiking of lNAc D1-MSNs and the spiking of lNAc-projecting VTA dopamine neurons, which argues against significant roles for local modulation of dopamine release^20,79^ or spiking adaptation in D1-MSNs in the TD error computation. We identify three categories of possible mechanisms: cell intrinsic mechanisms in dopamine neurons^80–82^, synaptic mechanisms in inputs to dopamine neurons^83,84^, and circuit mechanisms between D1-MSNs and dopamine neurons^27,83,85–87^. Cell intrinsic mechanisms could include depolarization block^82,88,89^ or D2 autoreceptor activation triggered by dopamine neuron spiking^90^. However, barring a spiking-independent mechanism, these mechanisms alone could not explain the pronounced suppression of action potential firing locked to the offset of pattern 3, which typically did not strongly activate spiking during the ramp-up phase (**Fig. 3i**). Next, synaptic mechanisms could include short-term synaptic depression (e.g. depletion of vesicle pools), which can create derivative-like responses in post-synaptic cells, wherein they respond proportionally to the percentage change of the input^84^. While short-term synaptic plasticity may play a role within a larger circuit computation^83^, a single depressing synapse cannot explain our finding that the dopamine response to lNAc D1-MSN stimulation plateaus at a level below baseline, which distinguishes TD error from a strict derivative. For these reasons, we hypothesize that circuit-level mechanisms play a major role in the temporal difference calculation. Importantly, although D1-MSNs are inhibitory neurons, activating them induced a burst of spikes in dopamine neurons (latency = 50-100 ms, **Extended Data Fig. 6c**), implying a disinhibitory mechanism. Disinhibitory circuit mechanisms could involve local inhibitory microcircuits within the VTA^83,85,91^ and neighboring substantia nigra pars reticulata (SNr), both of which receive innervation from lNAc D1-MSNs, or long-range multisynaptic basal ganglia circuits, such as from lNAc D1-MSNs to ventral pallidum (VP) to VTA/SNr^27,87^. Indeed, a recent study found that NAc D1-MSNs either (net) excite or (net) inhibit VTA dopamine neurons, depending on whether they project to the ventral midbrain (including VTA and SNr) or ventral pallidum^87^. If properly tuned in magnitude and timing, these oppositely signed responses could combine to form the temporal difference response reported here. The balance of these pathways would determine the temporal discount factor *γ*, thus grounding the TD learning algorithm’s sensitivity to time in a biological circuit mechanism (**Extended Data Fig. 10b,c**).

Altogether, our results show how TD computations are built into the circuitry of the basal ganglia, potentially explaining a wealth of neurophysiological observations, including ramping dopamine signals that possess a derivative-like property^5,19,20^. The same circuit mechanism may underlie temporal discounting at the level of single dopamine neurons^48,49^. While this shows that TD-like learning is a core property of the dopamine-striatum system when probed in isolation, natural behavior recruits additional circuits that may modulate or expand the functionality of the dopamine system. Therefore, the extent to which the LTI model proposed here can explain the contribution of dopamine and striatum to learning in more naturalistic contexts remains an important direction for future study.

---

**Extended Data Fig. 1:**
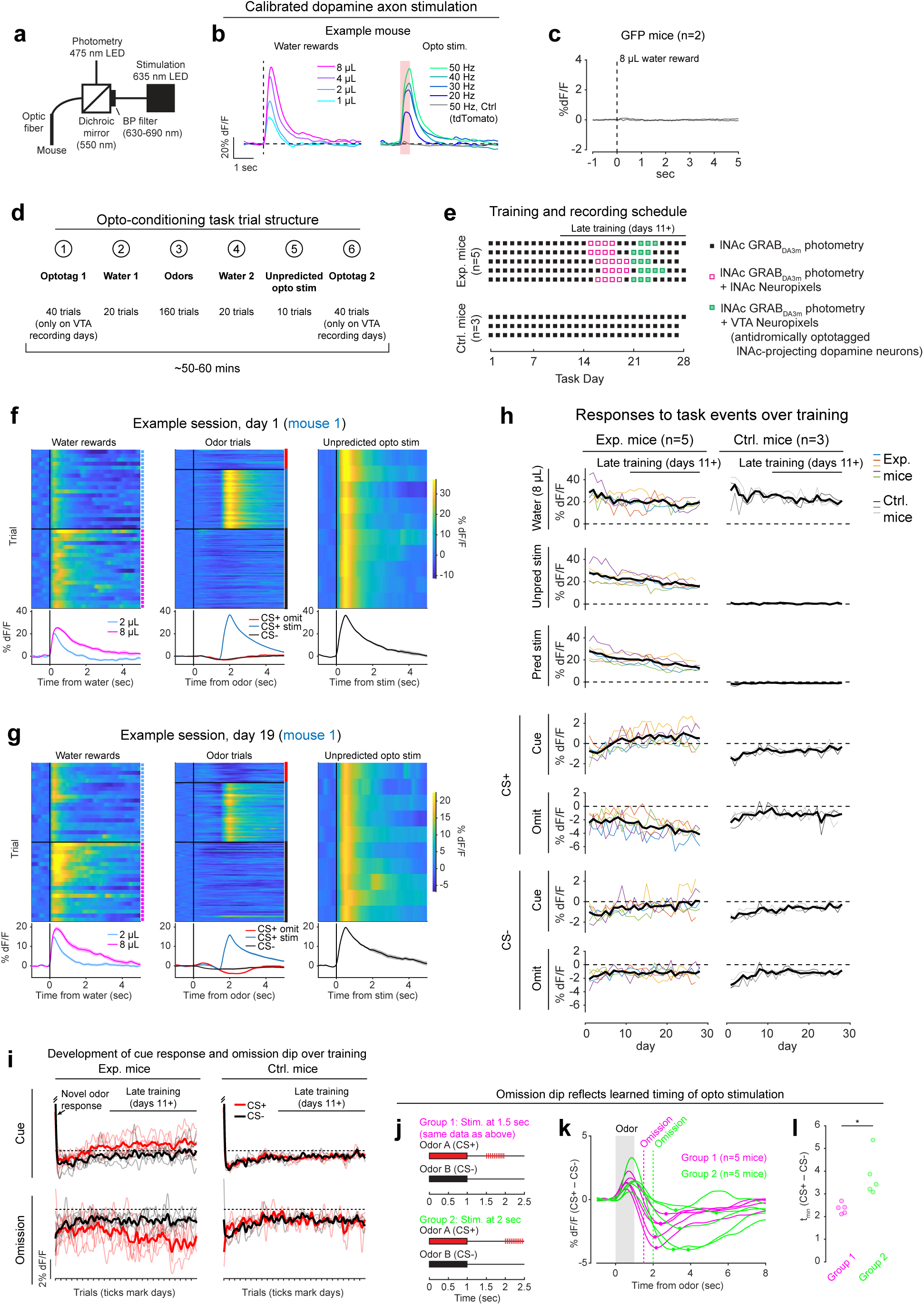
Calibration of opto-stimulation, opto-conditioning task design and training details, and opto-conditioning task responses over training. a) Schematic of experimental setup for optogenetic stimulation and photometry recording through the same optic fiber. b) Calibration of optical stimulation in an example mouse expressing ChrimsonR in VTA dopamine neurons and GRAB_DA3m_ in lNAc. *Left:* lNAc GRAB_DA3m_ signal in response to water rewards of various volumes (1, 2, 4, or 8 *μ*). *Right:* lNAc GRAB_DA3m_ signal in response to opto-stimulation pulse trains of various frequencies. All pulse trains lasted for 500 ms (e.g. 10 pulses at 20 Hz, 15 pulses at 30 Hz, etc.), individual pulses were 5 ms in duration, and red light power density was 40 mW/mm^2^ (5 mW through a 400 *μ* diameter fiber). The gray line shows the response to a 50 Hz pulse train in a different mouse expressing tdTomato instead of ChrimsonR in VTA dopamine neurons. Stimulation frequency was chosen to best match the response to 8 *μ* water reward (40 or 50 Hz, depending on the mouse). c) Photometry response to 8 *μ* water reward in mice expressing GFP instead of GRAB_DA3m_ in lNAc (*n* = 2 mice). We used 8 *μ* water delivery as a task event that elicited robust movements (licking) to check whether movement artifacts influence our photometry signals. The peak change in dF/F relative to baseline was < 0.15% for both mice, compared to > 60% in panel b, indicating that movement artifacts were negligible in our head-fixed setup. Because movement artifacts were negligible, and the isosbestic wavelength for GRAB_DA3m_ is unknown, we did not correct GRAB_DA3m_ recordings using an isosbestic channel. However, we did correct GCaMP recordings (**Fig. 2, Fig. 4**) using an isosbestic channel at *λ* = 415 nm, which is close to the known isosbestic wavelength for GCaMP sensors^92^. d) Trial structure of the opto-conditioning task. e) Training and recording schedule. Mice expressing ChrimsonR (Exp. Mice; *n* = 5) or tdTomato (Ctrl. Mice; *n* = 3) in VTA dopamine neurons were trained on the opto-conditioning task for up to 28 days. In exp. mice, Neuropixels recordings were performed during late training (defined as days 11+), in lNAc (on days 15-20) or VTA (on days 21-25). f) lNAc GRAB_DA3m_ signals in response to task events from an example Exp. Mouse on day 1 of training in the opto-conditioning task. Heatmaps show single trial responses, separated by trial type, and lines with shaded areas show mean responses ± SEM by trial type. Photometry signals were baseline-subtracted (the average response in the 1 second window before trial onset was subtracted before plotting the data). g) Same as panel f, but on day 19 of training. h) GRAB_DA3m_ responses to task events over training for Exp. Mice (left column) and Ctrl. Mice (right column) in VTA dopamine neurons. The period defined as “late training” (days 11+) is indicated by a black line above the first plot. Thin colored lines show individual mice, and the thick black line shows the average over mice. Event responses are defined as the mean photometry signal (dF/F, baseline subtracted) in the time window 0-1 second after event onset. i) The development of GRAB_DA3m_ responses to odors over training, at cue time (top row, 0-1 sec after odor onset) or omission time (bottom row, 1.5-2.5 sec after odor onset; only omission trials included for CS+), for exp. mice (left column) or ctrl. mice (right column). Single trial responses were averaged in the given time window, concatenated across days, and smoothed with a Gaussian filter (S.D. = 15 trials for all trial types except CS+ omission, which were 3x rarer and smoothed with an S.D. of 5 trials). Thin lines represent individual mice; thick lines show the mean over mice. j) To test whether the timing of the omission dip reflected the timing of opto-stimulation delivered during training, a second group of 5 mice (“Group 2”; green) was trained with opto-stimulation delivered at 2 seconds instead of 1.5 seconds. The original 5 exp. mice are shown for comparison (“Group 1”; magenta). k) Average difference in the GRAB_DA3m_ signal between CS+ omission trials and CS-in late training (days 11+) for Group 1 and Group 2 mice. The time of the omission dip was defined as the time of the minimum point of each trace, marked with a dot. l) Omission dip time was significantly later for Group 2 mice compared to Group 1 mice (mean minimum time ± SEM, Group 1 = 2.35 ± 0.10 sec, *n* = 5 mice; Group 2 = 3.79 ± 0.42 sec, *n* = 5 mice; *P =* 0.0102, unpaired t-test).

**Extended Data Fig. 2:**
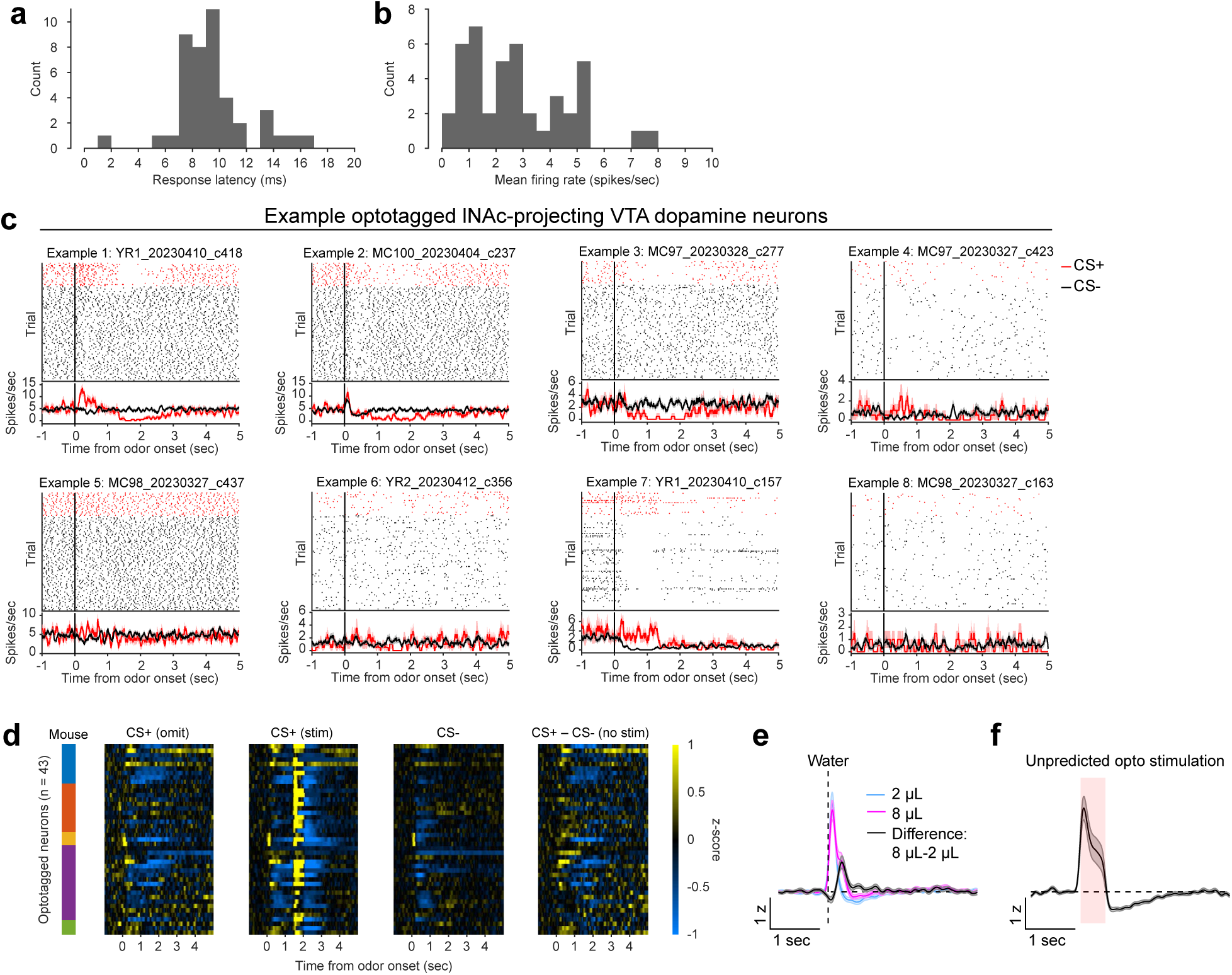
Additional detail on antidromic optotagging of lNAc-projecting VTA dopamine neurons during opto-conditioning. a) Histogram of spiking response latency to opto-stimulation pulse onset for optotagged lNAc-projecting VTA dopamine neurons (first significant bin ± SEM = 8.9 ± 0.4 ms, *n* = 43 neurons). b) Histogram of mean firing rate for optotagged lNAc-projecting VTA dopamine neurons (mean ± SEM = 2.8 ± 0.3 spikes/sec, *n* = 43 neurons). c) Example optotagged neuron responses to CS+ (omission trials, red) and CS-(black). Cell IDs are listed above each plot. The top panel of each plot shows spike rasters, split by trial type, and the bottom panel shows average firing rate by trial type (mean ± SEM). d) Heatmaps of all optotagged neurons’ z-scored responses to odors. Firing rates were z-scored to intertrial intervals (mean and standard deviation). The colored bars on the left indicate the mouse for each neuron. The four panels show: CS+ omission trials, CS+ stimulation trials (note the large response to opto-stimulation at 1.5 seconds, which exceeds the limits of the color scale; see panel f for response to unpredicted opto-stimulation), CS-trials, and the difference between CS+ and CS-trials. e) Z-scored response of optotagged neurons to unpredicted water delivery (mean ± SEM, *n* = 43 neurons). f) Z-scored response of optotagged neurons to unpredicted opto-stimulation (mean ± SEM, *n* = 43 neurons).

**Extended Data Fig. 3:**
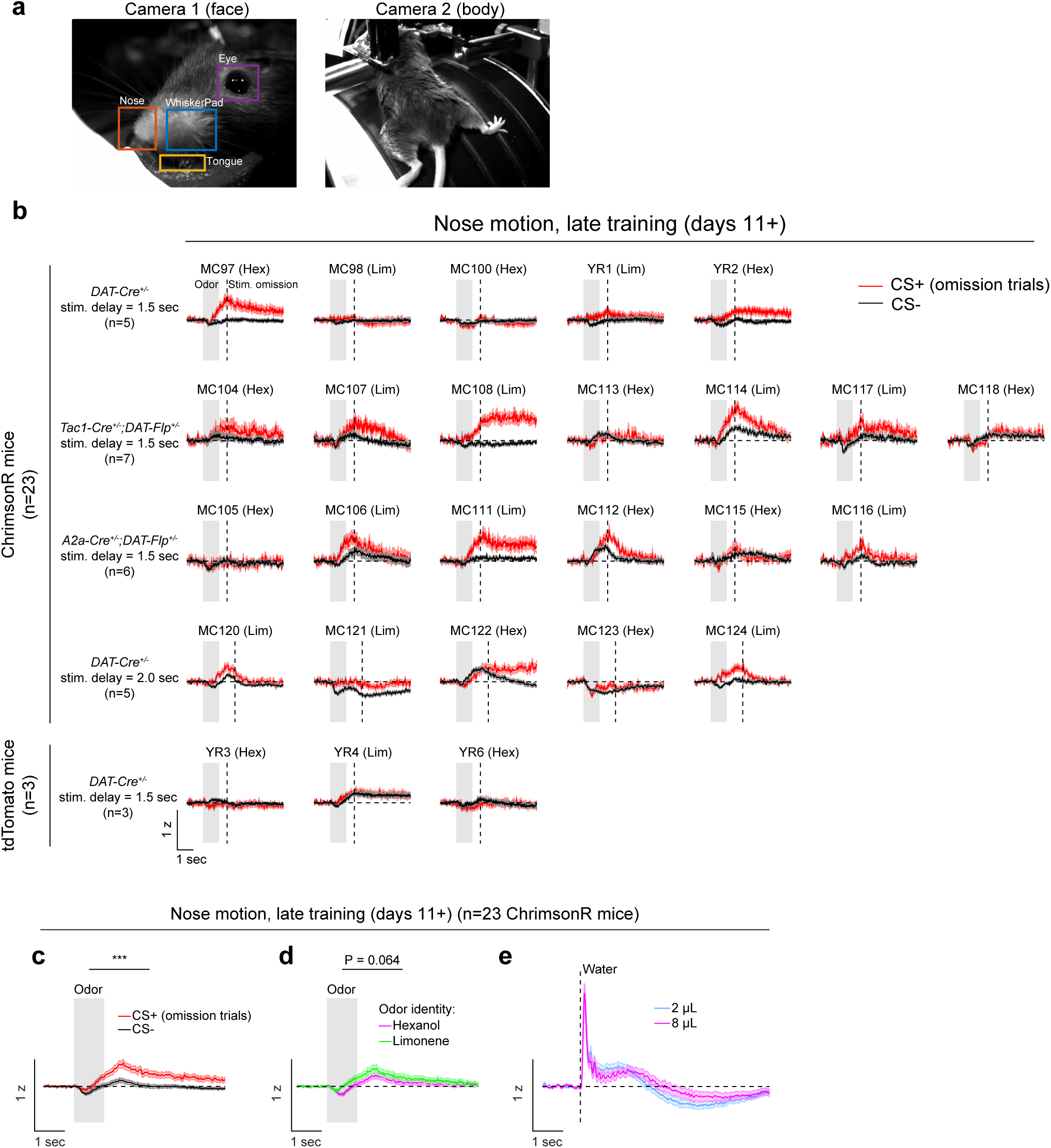
Opto-conditioning enhances facial movements following odor delivery. a) Example frames from the face camera (Camera 1; left) and body camera (Camera 2; right). To quantify behavioral responses to task events, motion energy (sum of absolute value of difference of subsequent frames) was computed within 5 ROIs, which were drawn manually for each video: WhiskerPad, Nose, Tongue, Eye (from Camera 1); and Body (the entire frame of Camera 2). b) Motion energy within the nose ROI, in late training of the opto-conditioning task (days 11+), in response to the CS+ (omission trials; red) or CS-(black) (mean ± SEM). Rows separate experimental conditions, which were combined for the analysis in panels c-e. Across the experiments of **Figures 1** and **2**, there were 23 ChrimsonR mice (top) and 3 tdTomato mice (bottom). The identity of the CS+ odor is listed above each plot (Hex = hexanol, Lim = limonene). c) Average motion energy within the nose ROI across ChrimsonR mice for CS+ omission trials (red) and CS-trials (black) (lines and shaded areas indicate mean ± SEM over *n* = 23 mice). Nose motion was greater following the CS+ compared to the CS-(mean ± SEM, 0.5 - 2.5 sec after odor onset, CS+ = 0.19 ± 0.04, CS-= 0.02 ± 0.03, *n* = 23 mice, *P =* 9×10^-6^, t-test). *** *P <* 0.001. d) Same as panel c, but split by odor identity (hexanol, magenta, and limonene, green). There was a marginal effect of odor identity, with greater nose motion to limonene compared to hexanol (mean ± SEM, 0.5 - 2.5 sec after odor onset, hexanol = 0.07 ± 0.03, limonene = 0.15 ± 0.04, *n* = 23 mice, *P =* 0.064, t-test). e) Motion energy within the nose ROI following water reward delivery, for comparison with odor-evoked nose movements.

**Extended Data Fig. 4:**
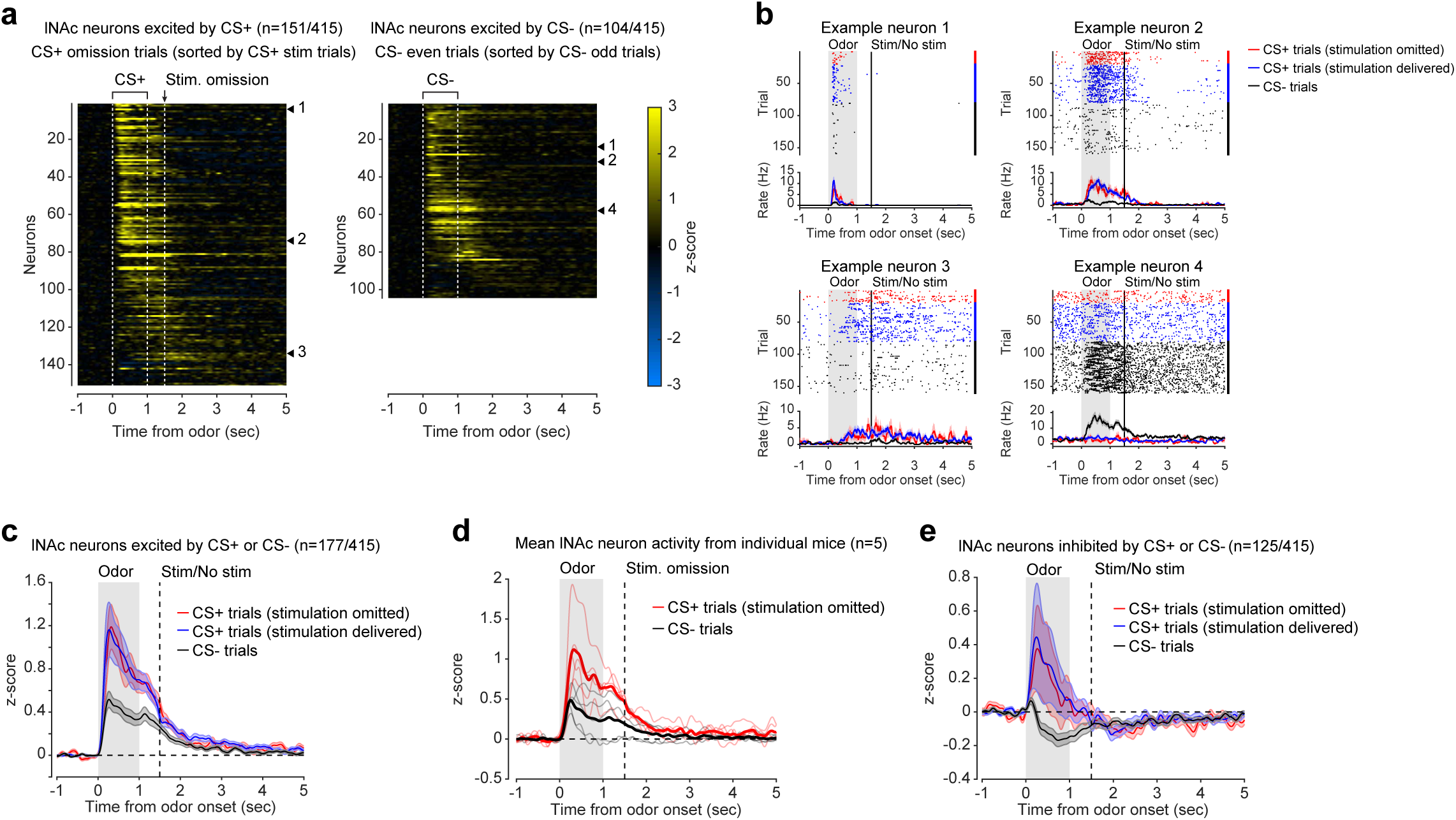
Additional detail on striatal Neuropixels recordings during opto-conditioning. a) Heatmaps of z-scored spiking responses of lNAc neurons to odors (CS+/CS-) in the opto-conditioning task. Only neurons identified as significantly excited by CS+ (left; *n* = 151/415 lNAc neurons) or by CS-(right; *n* = 104/415 lNAc neurons) are shown (Methods). For CS+, neurons were sorted based on the time of their peak response in CS+ stimulation trials, and responses on CS+ omission trials are shown (this was done due to the small number of omission trials and the lack of a difference between stimulation and omission trials; see e.g. panel c). For CS-, neurons were sorted based on the time of their peak response on odd CS-trials, and the response on even CS-trials is shown. The four example neurons shown in panel b are indicated. b) Spiking responses of four example neurons indicated in panel a. In this panel as well as panels c and e, trials are split into three groups: CS+ trials on which opto-stimulation was omitted (red), CS+ trials on which opto-stimulation was delivered (blue), and CS-trials (black). c) The average odor responses of lNAc neurons significantly excited by either CS+ or CS-in the opto-stimulation task (*n* = 177/415 lNAc neurons; same as **Fig. 2b** but including CS+ stimulation trials for comparison [blue trace]). Lines and shaded areas indicate mean ± SEM. d) The average odor responses of lNAc neurons significantly excited by either CS+ or CS-, averaged by mouse (*n* = 5 mice), on CS+ omission trials (red) or CS-trials (black). Thin red/black lines indicate individual mice; thick red/black lines show the average over mice. e) Same as panel c, but for neurons significantly inhibited by either CS+ or CS-(*n* = 125/415 lNAc neurons; Methods). Because CS+ responses were still positively enhanced relative to CS-responses in this group of neurons, we conclude that the primary effect of opto-stimulation was to positively enhance CS+ responses in a subset of neurons (as opposed to bidirectional positive/negative changes in CS+ response).

**Extended Data Fig. 5:**
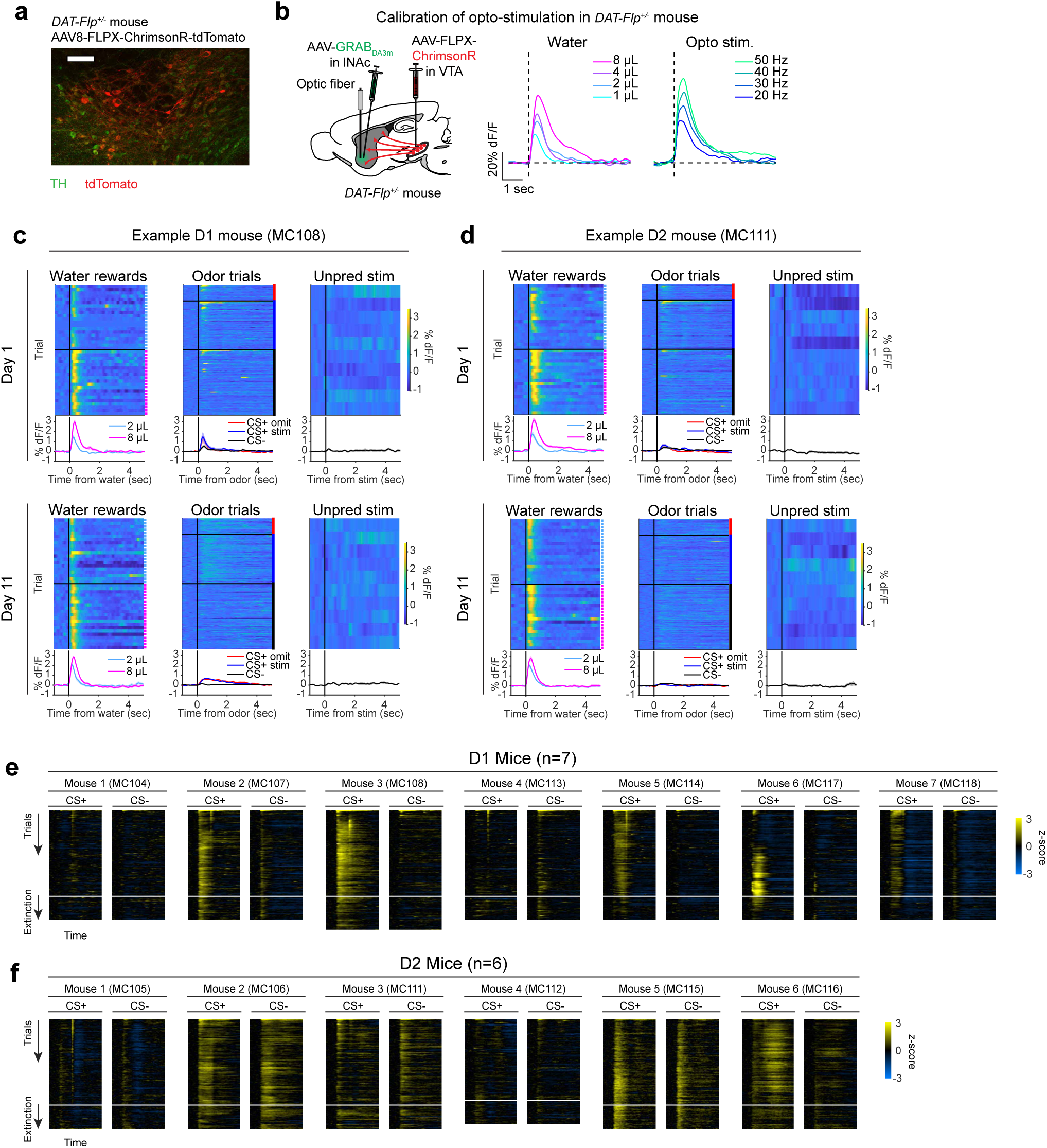
Validation and calibration of dopamine axon stimulation in *DAT-Flp^+/-^* mice, and development of D1- and D2-MSN responses to opto-conditioned odors over training. a) Confocal image of VTA in a *DAT-Flp^+/-^* mouse injected with AAV8-FLPX-ChrimsonR-tdTomato and stained for TH, showing co-expression of tdTomato and TH. Scale bar = 100 *μ*. b) Calibration of opto-stimulation to water reward in a *DAT-Flp^+/-^* mouse. Calibration could not be performed in the same mice as used for the experiment in **Fig. 2** because GCaMP8m was expressed in MSNs. Therefore, based on this panel, opto-stimulation frequency of 40 Hz was used for all mice, approximately matching an 8 *μ* water reward while remaining below the sensor’s ceiling. c) Example GCaMP8m photometry responses to task events (water rewards, odors, and unpredicted opto-stimulation) in a D1 mouse, on day 1 of training (top row) and day 11 of training (bottom row). Heatmaps show photometry responses on single trials, split by trial types. Traces and shaded areas show mean ± SEM for each trial type, averaged over trials. Note the emergence of a sustained photometry response to the CS+ odor, which was similar between trials on which opto-stimulation was delivered (blue) or omitted (red). d) Same as panel c, but in an example D2 mouse. Note that unlike the D1 mouse, photometry signals showed no difference between CS+ and CS-trials on day 11 of training. This was not due to a failure of sensor expression, as responses to water delivery and novel odors (first few trials) were observed. e) Heatmaps showing the development of odor responses in all individual D1 mice (*n* = 7) over training and extinction. The white line indicates the end of training and beginning of extinction. Single trials of each type (CS+ or CS-; CS+ omission and stimulation trials combined) were concatenated, forming a matrix of trials x time. To improve visualization, this matrix was smoothed over trials by applying a Gaussian filter to each column (corresponding to an individual time point within the trial) with SD = 15 trials. Note that some mice showed a photometry response to opto-stimulation which decays away over trials. Based on the experiment in **Fig. 2j**, we argue that this was a visual response to the red opto-stimulation light, not a response to dopamine release itself (no red masking light was used in this experiment). f) Same as panel e, but for D2 mice (*n* = 6).

**Extended Data Fig. 6:**
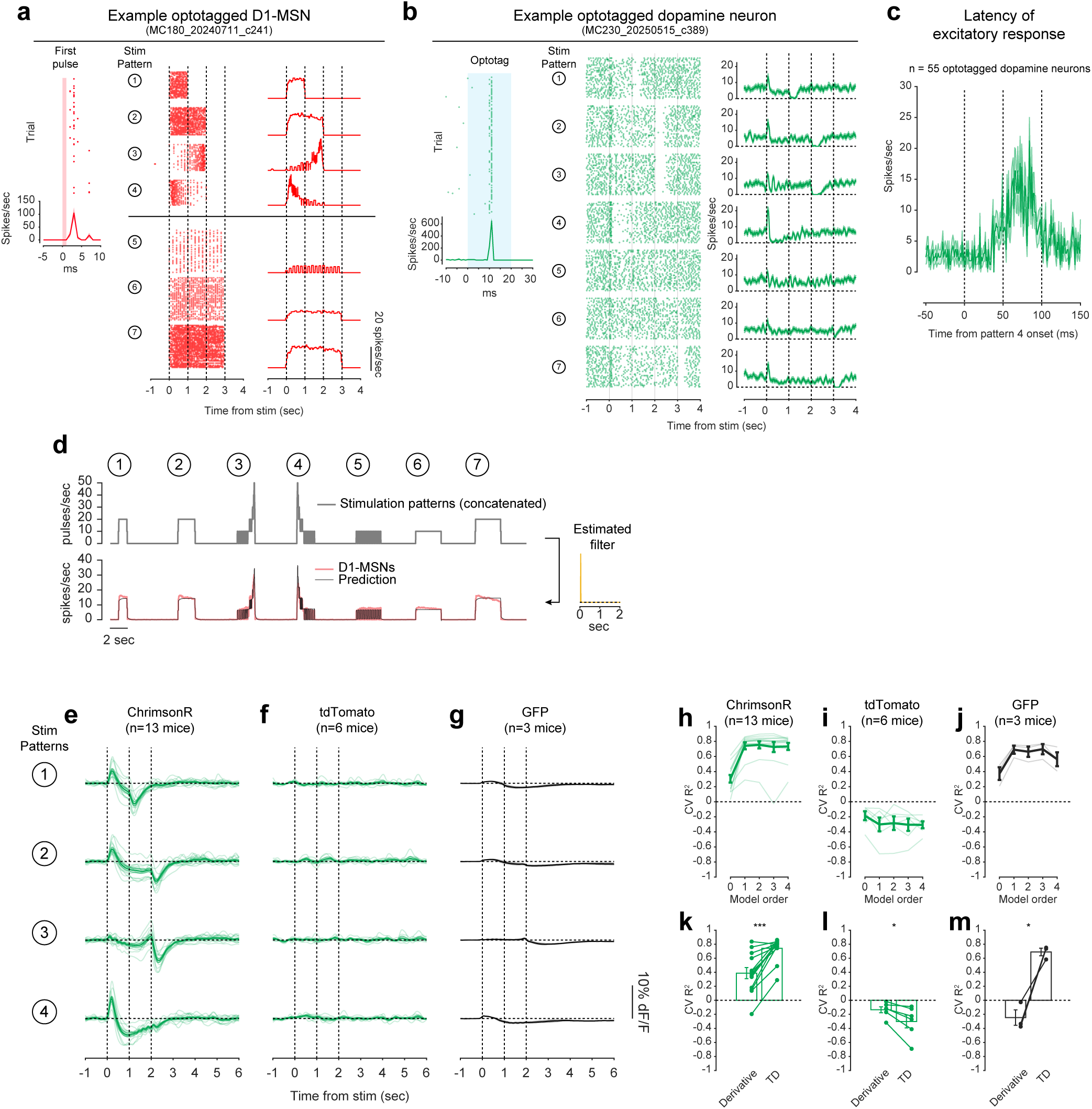
Additional detail on dopamine response to stimulation of lNAc D1-MSNs. a) Responses of an example single optotagged lNAc D1-MSN to opto-stimulation patterns of **Fig. 3**. *Left:* The response to the first pulse in the stimulus pulse train (all except patterns 3 and 4; used to identify optotagged neurons, Methods). *Middle:* Spike raster showing the response to each of the 7 stimulation patterns (same patterns as in **Fig. 3b**). *Right:* The firing rate in response to each of the 7 stimulation patterns. Lines and shaded area indicate mean ± SEM over trials. b) Responses of an example antidromically optotagged lNAc-projecting VTA dopamine neuron to opto-stimulation patterns of **Fig. 3**. *Left:* The response to the optotagging pulses (blue light; used to identify optotagged neurons, Methods). *Middle:* Spike raster showing the response to each of the 7 stimulation patterns (same patterns as in **Fig. 3b**). *Right:* The firing rate in response to each of the 7 stimulation patterns. Lines and shaded area indicate mean ± SEM over trials. c) Average PSTH of optotagged dopamine neurons (*n* = 55) aligned to the onset of pattern 4, which typically elicited a rapid burst of action potentials. Time scale is zoomed in to visualize the latency of the spiking response. Line and shaded area indicate mean ± SEM over neurons (*n* = 55). d) LTI model fit predicted D1-MSN response to the 7 stimulation patterns. *Top*: The 7 stimulation patterns, concatenated. *Bottom*: Average lNAc D1-MSN responses to the 7 stimulation patterns (red; averaged over *n* = 2 mice) and the LTI model fit (black) predicting D1-MSN response (output) from stimulation patterns (input) (cross-validated [CV] R^2^ = 0.95 Methods). *Right*: Estimated filter of the LTI model. Note that the filter is approximately a delta function, which scales the input to produce the output. e) The GRAB_DA3m_ photometry response to opto-stimulation patterns 1-4 in mice prepared as in **Fig. 3a**, with optogenetic stimulation of lNAc D1-MSNs and photometric recording of lNAc dopamine release, but without simultaneous Neuropixels recording of optotagged lNAc D1-MSNs. The stimulation parameters differed: here, we used 5 ms instead of 1 ms pulses, and power was reduced 5x from 40 mW/mm^2^ to 8 mW/mm^2^ (1 mW through a 400 µm diameter fiber) (same parameters used in panels e-g). The TD error-like response patterns were the same as in **Fig. 3c** and **Fig. 3e**, with positive/negative transients locked to the onset/offset of constant stimulation and a negative plateau before offset; a strong positive response to pattern 4 (ramp down) but not pattern 3 (ramp up); and a negative response to the offset of pattern 3. Thick line and shaded area indicate mean ± SEM over mice (*n* = 13). Thin lines indicate individual mice. f) Same as panel e but for mice with tdTomato expressed instead of ChrimsonR in lNAc D1-MSNs. No significant responses to optical stimulation were observed, ruling out optical crosstalk and visual responses. g) Same as panel e, but for mice with GFP expressed instead of GRAB_DA3m_ in VTA axons in lNAc. In contrast to the no-opsin (tdTomato) control, small fluctuations in the GFP signal were observed following optical stimulation of D1-MSNs, possibly due to hemodynamic responses to optogenetic stimulation. h) Same as **Fig. 3f**, but for the mice in panel e (*n* = 13). In panels h-j, thin lines indicate individual mice and thick lines and error bars indicate mean ± SEM over mice. In panels h-m, LTI models were fit predicting dopamine responses directly from stimulation patterns, as D1-MSN activity was not recorded in these mice. i) Same as panel h, but for tdTomato control mice (*n* = 6). j) Same as panel h, but for GFP control mice (*n* = 3). k) Same as **Fig. 3g**, but for the mice in panel e (*n* = 13). In panels k-m, lines indicate individual mice and bars/error bars indicate mean ± SEM over mice. *** *P =* 6.8 × 10^-5^, paired t-test. l) Same as panel k, but for tdTomato control mice (*n* = 6). * *P =* 0.022, paired t-test. m) Same as panel k, but for GFP control mice (*n* = 3). * *P =* 0.029, paired t-test.

**Extended Data Fig. 7:**
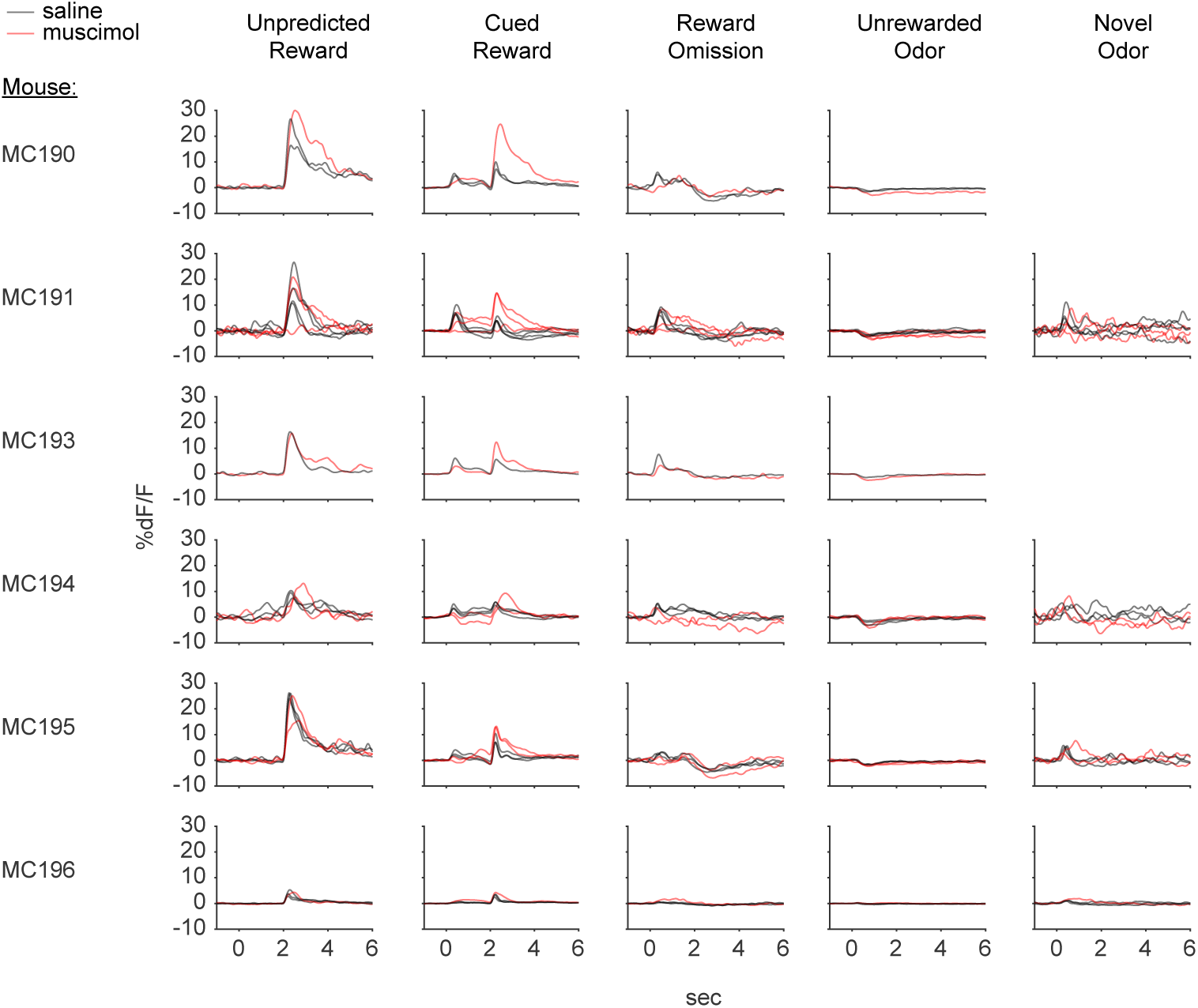
Additional detail on VTA dopamine neuron calcium activity during injection of muscimol or saline into lNAc (related to Fig. 4) Average VTA dopamine neuron calcium response aligned to task events for each session in the data set (*n* = 26 session from 6 mice). Rows indicate mice, columns indicate trial types, lines show the average response for that trial type within an individual session, and colors indicate saline (black) versus muscimol (red) injection. Note that novel odors were only delivered for 4 of 6 mice (21/26 sessions).

**Extended Data Fig. 8:**
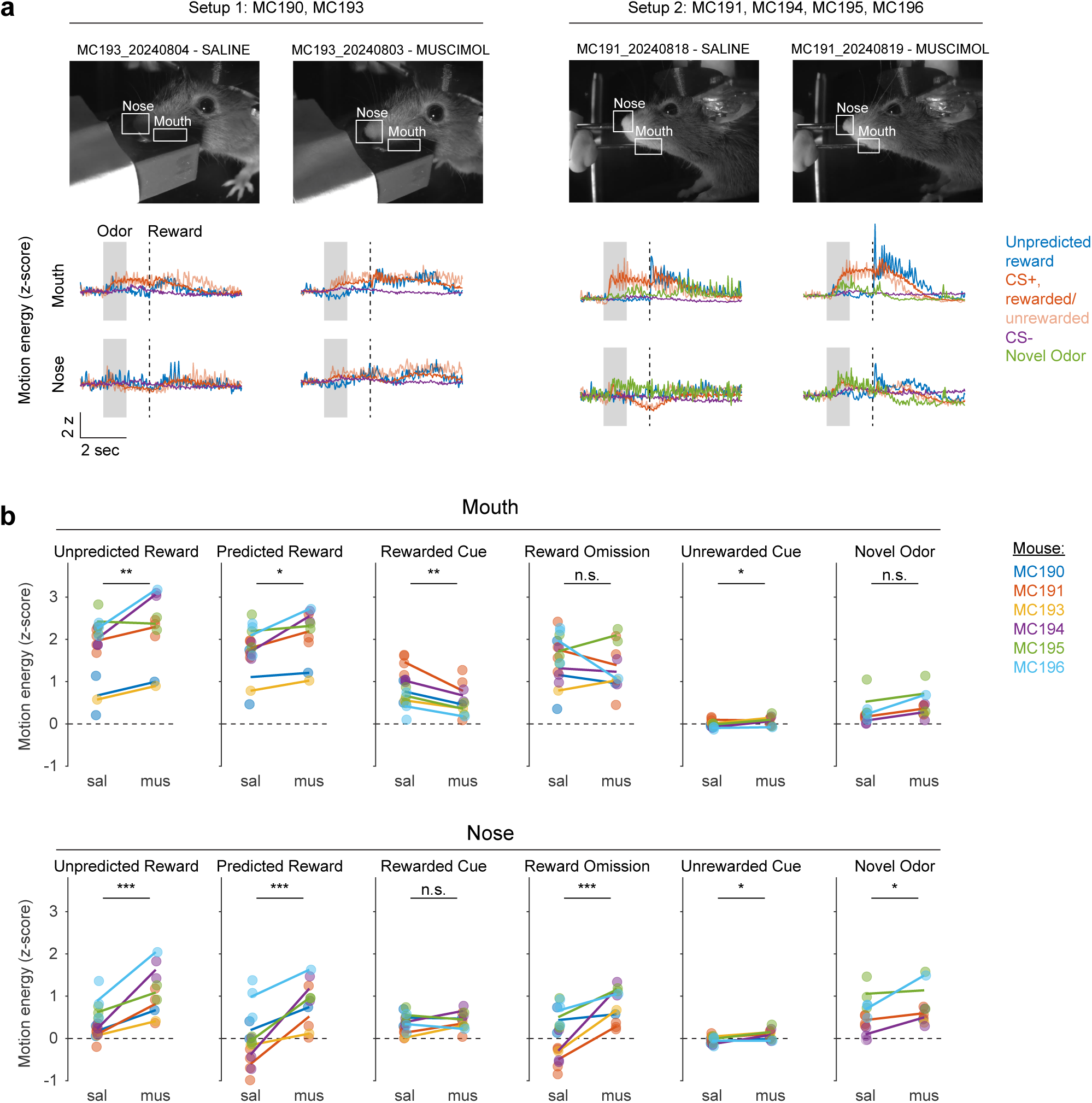
Facial movements in classical conditioning task with muscimol versus saline in lNAc. a) Example video frames (top) and task-aligned motion energy traces (bottom) from the muscimol experiment (**Fig. 4**). Two different odor/water delivery setups were used in the experiment: Setup 1 (left) was used for mice MC190 and MC193, and Setup 2 (right) was used for mice MC191, MC194, MC195, and MC196. Example frames and average motion energy traces are shown for an example saline day and muscimol day for each setup. ROIs were manually drawn around the nose and mouth, motion energy (sum of absolute value of the difference between subsequent frames) was computed within each ROI, and z-scored across the entire session. Novel odors were not delivered in every session, but an example pair of sessions is shown in which novel odors were also delivered (green traces, right). b) Average z-scored motion energy in the mouth ROI (top) and nose ROI (bottom) in 1-second windows following task events during saline (sal) and muscimol (mus) sessions. Lines indicate average over sessions for each mouse and points indicate individual sessions; colors indicate mice. Significance stars indicate significance of treatment (saline versus muscimol) in a linear mixed effects model with a fixed effect of treatment and random effect per mouse. n.s. Not Significant, * *P <* 0.01, ** *P <* 0.001, *** *P <* 0.0001.

**Extended Data Fig. 9:**
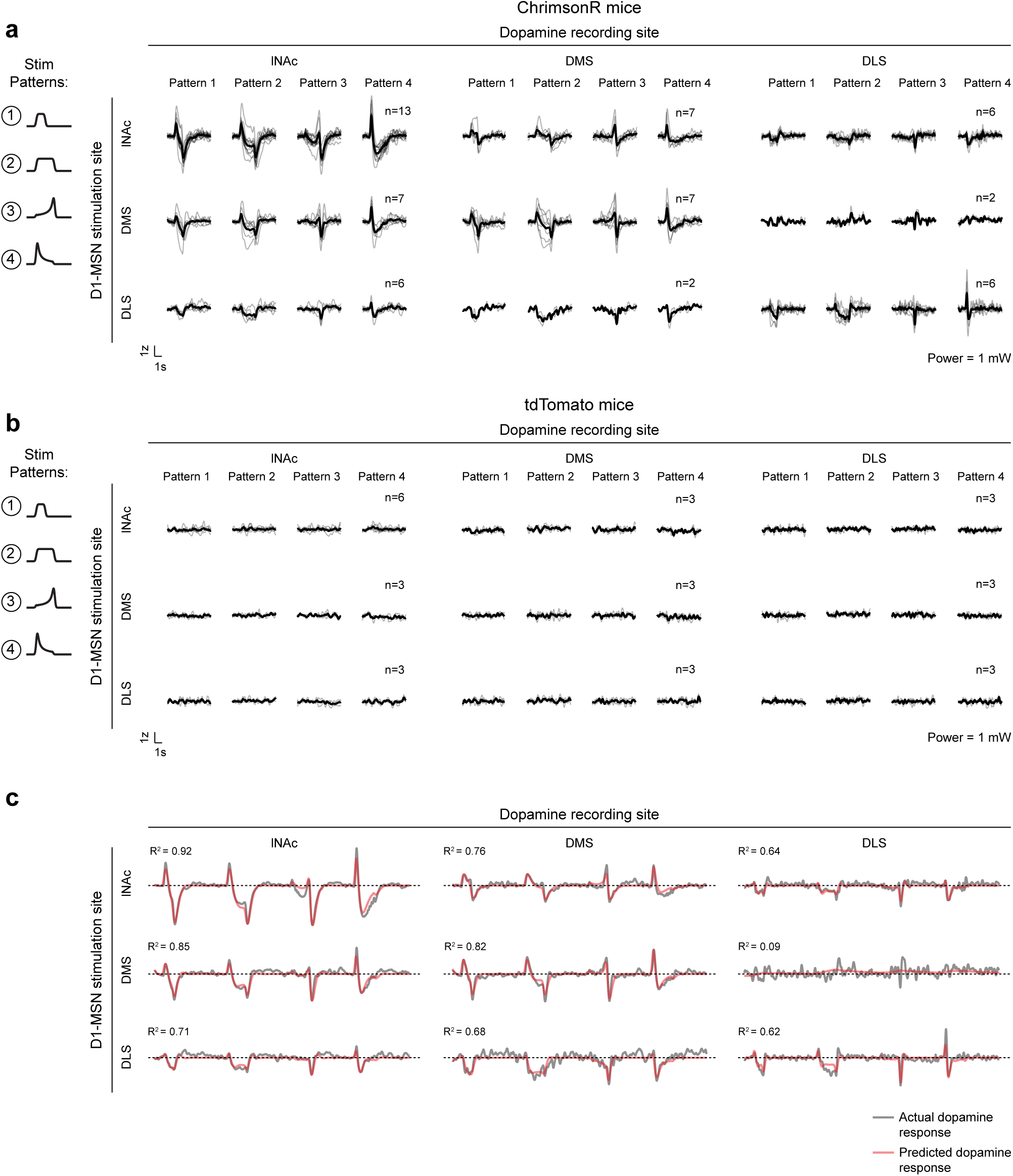
Additional detail on dopamine responses to stimulation of D1-MSNs in multiple striatal subregions (lNAc, DMS, DLS) a) In this experiment, only the first 4 of the 7 stimulation patterns were used. Photometry responses to the 4 stimulation patterns (schematized to the left of the figure) for each pair of D1-MSN stimulation site (rows) and dopamine sensor photometry recording site (columns), as in **Fig. 5b** but showing the response to all 4 patterns. As in **Fig. 3** and **Fig. 5**, dopamine photometry recordings were made by expressing GRAB_DA3m_ non-specifically in VTA neurons and recording photometry signals from VTA axons in the given striatal recording site. Black lines show the mean response across animals and gray lines show individual animals. The number of animals for each pair of stimulation and recording sites is listed above the response to Pattern 4. Red light stimulation power was 1mW through a 400 *μ* diameter optical fiber (8 mW/mm^2^). b) Same as panel a, but for control mice expressing tdTomato instead of ChrimsonR in D1-MSNs. No significant responses were observed, indicating that optical crosstalk was successfully eliminated, and that the red stimulation light alone was not sufficient to drive visual dopamine responses (red masking lamps were used in these experiments to mask red stimulation light). c) For each pair of D1-MSN stimulation site (rows) and dopamine sensor photometry recording site (columns), the actual response to the four stimulation patterns of panels a and b (black; averaged over mice) and the predicted response of an LTI model with 2 poles and 1 zero (red; fit using the function *tfest* in MATLAB, from the Systems Identification Toolbox, as in **Fig. 3, Fig. 5**). The *R*^2^ between the data and model fit is listed above each plot, corresponding to the given pair of stimulation and recording sites.

**Extended Data Fig. 10:**
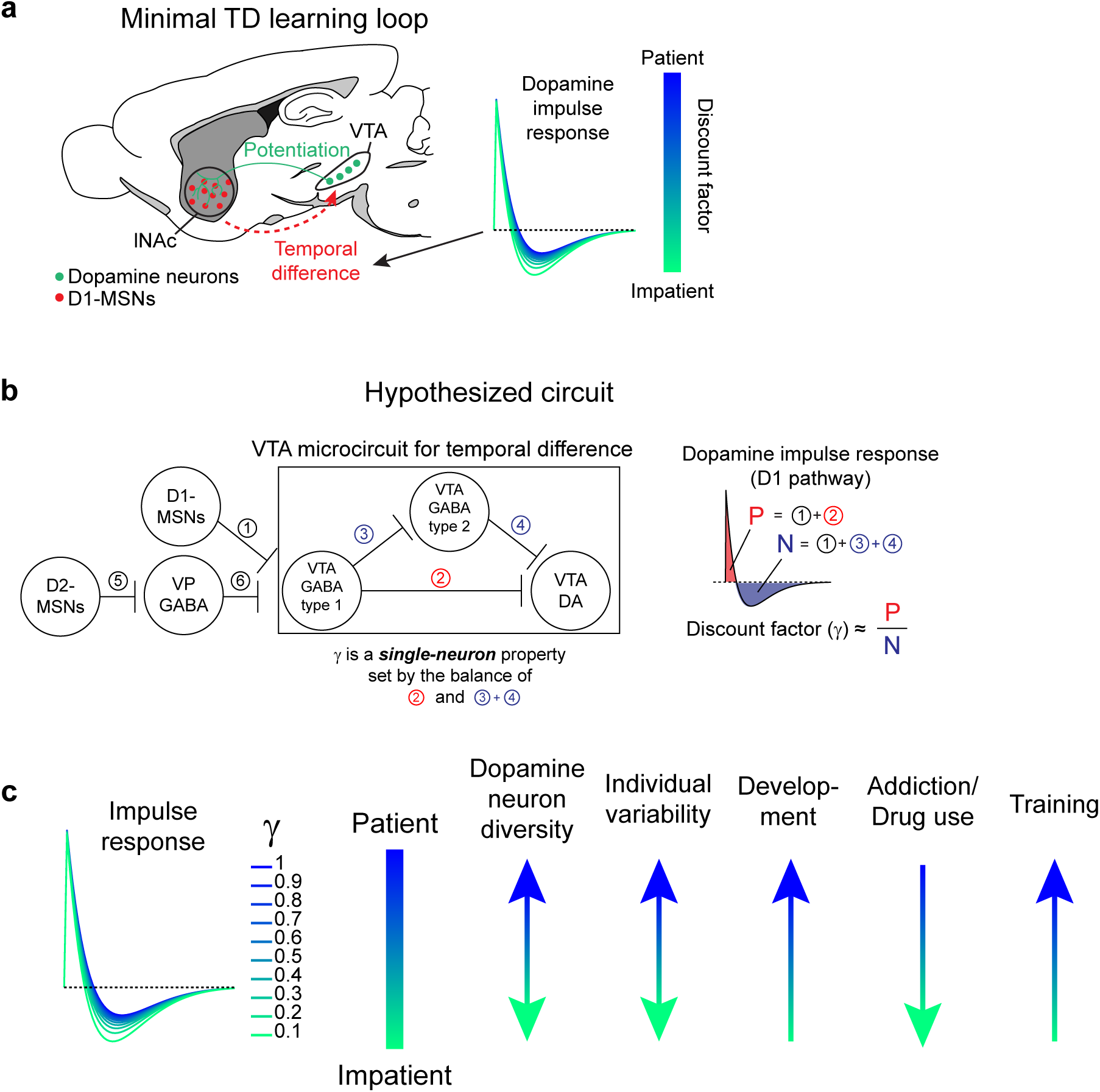
Summary of findings and hypotheses for future work. a) Minimal TD learning loop, based on the experiments of **Fig. 1-3**. Calibrated dopamine release in lNAc potentiates the response of lNAc D1-MSNs to preceding sensory stimuli (green line, indicating dopaminergic axons projecting from VTA to lNAc); in turn, lNAc D1-MSNs generate TD error-like signals in lNAc-projecting VTA dopamine neurons via a temporal difference calculation which occurs between lNAc and VTA (red dashed line, indicating a net circuit effect). The temporal difference calculation can be captured by convolution with a biphasic linear filter, whose shape sets the temporal discount factor of the TD algorithm. b) Schematic of a possible neural circuit computing a temporal difference involving inhibitory and disinhibitory microcircuits within the VTA^83,85,93^. VP: ventral pallidum. c) Potential connections of the shape of the dopamine impulse response to temporal discounting at the neuronal and behavioral levels.

## Methods

### Experimental procedures

#### Animals

A total of 74 mice (39 male, 35 female, aged 2-12 months at time of surgery) were used in the experiments. Mice were housed on a 12 hr / 12 hr dark/light cycle and experiments were performed in the dark phase. All procedures were performed in accordance with the National Institutes of Health Guide for the Care and Use of Laboratory Animals and approved by the Harvard Institutional Animal Care and Use Committee.

#### Surgeries

All surgical procedures were performed in aseptic conditions. Mice were anesthetized using isoflurane (4% induction, 1-2% maintenance) and given an intraperitoneal injection of buprenorphine HCl (0.1 mg/kg) for analgesia. They were then head-fixed within a stereotactic surgical setup (Narishige). The hair was removed, the scalp was cleaned with 70% ethanol then 10% povidone-iodine, and a local anesthetic (2% lidocaine, 0.05 mL) was injected under the scalp. The scalp and periosteum were removed and the skull was dried and scored with a scalpel blade to ensure good adhesive binding. The skull was leveled by ensuring bregma and lambda were at the same DV position; no ML leveling was performed. Fiducial marks were made above the target sites (for virus injections, fiber implants, electrophysiology recording, or muscimol injection) using a fine-tipped pen. The following coordinates were used, with all distances in mm (AP and ML measured relative to bregma and DV measured from the surface of the brain):

VTA, virus injection (**Fig. 1-3, 5**): −3.0 AP, 0.6 ML, 4.3→4.2→4.1 DV
VTA, fiber implant (**Fig. 4**): −3.0 AP, 0.6 ML, 4.2 DV
SNc, virus injection (**Fig. 5**): −3.0 AP, 1.6 ML, 4.2→4.1→4.0 DV
lNAc virus injection (**Fig. 1-3, 5**): 1.3 AP, 1.8 ML, 4.1 DV
lNAc fiber implant (**Fig. 2, 3, 5**): 1.3 AP, 1.8 ML, 4.0 DV
lNAc angled fiber implant (**Fig. 1-3**): 1.3 AP, −1.1 ML, 4.75 DV, angled 35° in coronal plane away from midline
DMS, virus injection (**Fig. 5**): 0.5 AP, 0.5-0.7 ML, 2.7→2.6 DV, angled 15° in coronal plane towards midline
DMS, fiber implant (**Fig. 5**): 0.5 AP, 0.5-0.7 ML, 2.5 DV, angled 15° in coronal plane towards midline DLS, virus injection (**Fig. 5**): −0.3 AP, 2.8 ML, 2.6 DV, angled 15° in sagittal plane towards posterior
DLS, mirrored fiber implant (**Fig. 5**): −0.3 AP, 2.6 AP, 2.8 DV, angled 15° in sagittal plane towards posterior, mirror facing towards lateral side

The exact combinations of viruses, fibers, and target sites depended on the experiment and are given in the corresponding sections. Craniotomies were made above virus injection and fiber implant target sites using a high-speed dental drill. Virus injections were performed using glass pipettes (Drummond, pulled using Narishige PC-10) with an oil hydraulic microinjector (Narishige MO-10). All virus injections were 300 nL volume with viral titer ranging from 5 × 10^12^ to 2 × 10^13^ gc/mL. The pipette was slowly withdrawn 10 minutes after completing the injection. Optic fiber cannulae were scored using the dental drill (to improve adhesive binding), slowly implanted using a stereotaxic cannula holder (SCH_1.25, Doric Lenses), and temporarily held in place using a small amount of superglue (n-butyl cyanoacrylate) and accelerant. After fiber implants, a custom titanium headplate was attached to the skull with dental cement (Metabond, Parkell) containing a small amount of charcoal powder to reduce light leakage. The cement also served to reinforce the optic fiber implants. For electrophysiology and muscimol experiments, a 3D-printed plastic well was glued to the headplate with epoxy or superglue, to hold saline during recordings or injections. For electrophysiology experiments, a ground pin coated with a lead-tin alloy (Mill-Max) was implanted above contralateral cortex. Apart from ground pin implants, all surgical procedures (virus injections, fiber implants, craniotomies) were done in the left side of the brain.

Following surgery, mice recovered on a heating pad, and were given ketoprofen (20 mg/kg, injected SC) and 0.5 mL of sterile saline (injected IP). Mice were then singly housed and post-operative analgesia (ketoprofen, 20 mg/kg) was given twice daily for 48 hours. In experiments using viruses, we waited at least 3 weeks for the virus to express before starting experiments.

### Behavioral training

All behavioral experiments took place inside custom-built, partially sound-proofed, enclosed behavioral setups in which head-fixed mice ran freely on a cylindrical running wheel. White noise machines outside the setups were used to mask background noise. Except for opto-stim experiments using red masking lamps (**Fig. 2j, Fig. 3, Fig. 5**; see *Red masking lamp* below), experiments were run in the dark, with infrared lamps illuminating the mouse for face and body videos. Behavioral tasks were controlled using a Bpod state machine (1027, Sanworks) and accessories (1004: Port Interface, 1020: Optical Lickometer, 1038: Analog Output Module, 1039: Valve Module; Sanworks) with custom software written in MATLAB (R2021b) based on functionality in the Bpod MATLAB Software Repository (github.com/Sanworks/Bpod_Gen2).

#### Water deprivation

Mice were water-deprived starting at least 7 days after surgery. After this, mice were weighed daily and given water (1-2 mL) to maintain their bodyweight above 85% of baseline throughout the experiment.

#### Water delivery and lick detection

Water droplets were delivered to head-fixed mice through a spout attached to an optical lickometer (1020, Sanworks), which detected licks as breaks of an infrared beam. Water delivery was controlled by a Bpod port interface (1004, Sanworks) and valve (LHDA1231115H, The Lee Company). Water droplet volume was calibrated using a built-in Bpod protocol. In some experiments, to improve visibility of the face, the beam-break lickometer was removed and licks were detected directly from face videos in post-processing (e.g. **Extended Data Fig. 8a**, right).

#### Habituation to recording setups

All mice went through the same habituation protocol before starting experiments: 1 day of handling by experimenter, 1 day of freely exploring the head-fixation setup while receiving water from a syringe (administered by experimenter), 1 day of head-fixation while receiving water from a syringe (administered by experimenter), followed by several days of head-fixation with water droplets administered from the same valve/spout used in experiments. Thus, when mice began experiments, their weights had stabilized, and they were well habituated to handling by the experimenter, head-fixation, and receiving water droplets within the recording setup.

#### Odor preparation and delivery

Pure odors (1-hexanol, (S)-(−)-limonene, isoamyl acetate, p-cymene, ethyl butyrate, (+)-carvone, (+/-)-citronellal, eugenol, 1,4-cineole, L-fenchone, α-ionone; Sigma-Aldrich) were diluted to 10% in mineral oil (Sigma-Aldrich). Each day, 30 *μ* of each odor solution used in the current experiment was placed on a syringe filter (Whatman Puradisc 6823-1327; 13 mm diameter, 2.7 *μ* pore size) and the filters were attached to a custom odor manifold made of polyetheretherketone (PEEK). Filtered air (0.9 L/min) was constantly delivered to the mouse’s nose. A system of valves (LHDA1221111H, The Lee Company, mounted in a manifold, LFMX0510528B, The Lee Company), controlled by the Bpod state machine (1027, Sanworks) with an accessory valve module (1015, Sanworks), added 0.1 L/min to this air stream, for a total flow of 1 L/min. By default, this added air passed through a filter containing only mineral oil (“blank”). To deliver an odor, the valves switched the additional air flow from the blank filter to the desired odor filter. A vacuum line inside the recording setup provided air circulation to remove odors. In all experiments, odor identities assigned to CS+/CS-were counterbalanced across mice.

#### Opto-conditioning task

After standard habituation, mice started training on an odor-based opto-conditioning task (**Fig. 1, Fig. 2**). The two odors used were 1-hexanol and limonene. One odor was associated with opto-stimulation of dopamine axons in lNAc on 75% of trials (CS+) and the other was associated with nothing (CS-). Dopamine axon stimulation was calibrated to a natural reward (8 *μ* water) for each mouse (**Fig. 1e, Extended Data Fig. 1b, Extended Data Fig. 5b**; see **Fiber photometry with optogenetics** for details). Odors were delivered for 1 second followed by a 0.5 second trace period (**Fig. 1, Fig. 2**) or 1 second trace period (**Extended Data Fig. 1j-l**) before opto-stimulation. Inter-trial intervals (ITIs) were drawn from a truncated exponential distribution with mean 4 seconds and maximum 12 seconds and added to a fixed 8.5 second interval for a total ITI of 8.5-20.5 seconds with mean 12.5 seconds. 80 trials of each type (CS+ and CS-) were delivered, pseudo-randomly interleaved, for a total of 60 CS+ stimulated trials, 20 CS+ omission trials, and 80 CS-trials. Immediately before and after the odor trials, we delivered blocks of water rewards, as a comparison point for the signals generated by odors and opto-stimulation. These blocks of water rewards consisted of 16-20 trials, split evenly between 2 *μ* and 8 *μ* droplets pseudo-randomly interleaved, with the same ITI distribution as the odor trials. At the very end of the session (after the second block of water rewards), 10 unpredicted opto-stimulation trials were delivered, with the same ITI distribution, to compare predicted and unpredicted opto-stimulation responses. Mice were trained in this task for up to 30 days, with photometry recording each day, and electrophysiology recordings on some days in late training (days 11+) (see **Fiber photometry with optogenetics** and **Electrophysiology recording and spike sorting** for details).

#### Classical conditioning task with water rewards

After standard habituation, mice started training on an odor-based classical conditioning task with water rewards (**Fig. 4**). The two odors used were isoamyl acetate and p-cymene. In the first phase, odors were paired with 100% reward delivery (4 *μ* water) or nothing. Odors were delivered for 1 second followed by a 1 second trace period. Inter-trial intervals (ITIs) were drawn from a truncated exponential distribution with mean 4 seconds and maximum 12 seconds and added to a fixed 8.5 second interval for a total ITI of 8.5-20.5 seconds with mean 12.5 seconds. 80 trials of each type were delivered each day for 2-6 days. One mouse (MC190) started training with probabilistic reward delivery (80-90%) for 7 days before switching to 100% reward delivery for 3 days. After this shaping phase, mice were switched to the full task, in which the CS+ switched from 100% to 90% rewarded (in 1/10 trials, reward was omitted), and rare unpredicted rewards were delivered (1 in 21 trials). After training in the full task for 1-3 days, a craniotomy was performed over lNAc and muscimol or saline was injected into lNAc before the task each day for up to 8 days (see **Muscimol injections**). In some muscimol/saline sessions, we also delivered novel odors (5 or 10 trials total), either interleaved with other trial types, or at the end of the recording session. A different novel odor was used each day. The novel odors used were: 1-hexanol, (S)-(−)-limonene, ethyl butyrate, (+/-)-citronellal, eugenol, 1,4-cineole, L-fenchone, α-ionone.

#### Red masking lamp

In some experiments, including all experiments in which D1-MSNs were stimulated while recording dopamine signals (**Fig. 3, Fig. 5**) as well as the control experiment in **Fig. 2j**, the recording setups were illuminated with red light to mask visual input from red optogenetic stimulation light. Red light illumination came from two lamps with red light bulbs pointed directly at the mouse’s eyes. This eliminated visual dopamine responses to red light (**Fig. 2j**) so that optogenetic responses could be better isolated.

### Face and body videos

In all experiments, face and body videos were recorded using infrared cameras (Blackfly BFLY-U3-03S2M, FLIR), running at least 30 frames per second, with custom software in Bonsai (https://bonsai-rx.org). Videos were synchronized to other data streams using TTL pulses output by BPod at the start of each trial, detected by cameras’ GPIO lines and saved to file for each video frame.

### Fiber photometry with optogenetics

Photometry recordings were performed using a commercial photometry imaging system (Bundle-imaging Fiber Photometry System, Doric Lenses). A branching bundle patch cord (BBP(3)_400/440/900-0.37_1m_FCM-3xFC_LAF, Doric Lenses) was imaged by a CMOS camera (Sony IMX249LLJ) through an objective. The branching patch cord split into three branches, which could be used to image up to three sites simultaneously. A blue excitation LED (470 nm, 50 *μ* power at tip of each 400 *μ* diameter patch cord, 20 ms illumination at 20 Hz) was used to collect GRAB_DA3m_ (**Fig. 1, 3, 5**), GCaMP8m (**Fig. 2**), or GCaMP6f (**Fig. 4**) signals. A purple excitation LED (415 nm, 50 *μ* power at tip of patch cord, 20 ms illumination at 20 Hz, interleaved with blue excitation pulses) was used to collect isosbestic signals, but these were only used to correct GCaMP signals as the isosbestic wavelength for GRAB_DA3m_ is not known. GRAB_DA3m_ recordings were not corrected by a control channel. Instead, control GFP mice were used to rule out movement artifacts, which were minimal in our head-fixed setup (**Extended Data Fig. 1c**). In experiments in which GRAB_DA3m_ and Neuropixels recordings were made at the same site (**Fig. 2, Fig. 3**), blue excitation light power was reduced to 20 *μ* and purple excitation light was turned off to minimize light artifacts in the Neuropixels recording.

To perform photometry and optogenetics through the same optic fiber implant, red excitation light was independently coupled to each of the three patch cord branches using a fluorescence minicube (FMC3_(390-540)_(560-1000)_S, Doric Lenses). Each minicube contained a dichroic mirror which reflected blue/green photometry light and passed red optogenetic excitation light (cutoff wavelength ≈ 550 nm). Each minicube had three FC connectorized ports, which were connected to: (1) one branch of the bundle-branching patch cord, the other end of which connected to the Doric photometry setup, (2) a 635 nm red LED (CLED_635, Doric Lenses), and (3) a patch cord (MFP_400/440/110-0.37_2m_FC-ZH1.25(F)_LAF, Doric Lenses) connecting to the corresponding fiber implant on the animal. To further reduce optical crosstalk, the output of the red LED was filtered by a band-pass filter (Semrock FF01-630/92, transmitting 580-680 nm) installed in the corresponding minicube port. Each LED’s output was triggered by TTL pulses from the Bpod state machine, which controlled the behavioral task. With this setup, red optogenetic excitation light could be independently delivered to each of up to three optic fiber implants while simultaneously collecting photometry recordings. The lack of optical crosstalk was confirmed using recordings in tdTomato control animals (**Extended Data Fig. 1b**).

### Electrophysiological recording

Spiking data were collected using Neuropixels 1.0 single shank probes^94^ or Neuropixels 2.0 four shank probes^95^. Neuropixels data were acquired via a National Instruments PXI chassis (NI PXIe-1061, with NI PXIe-8381 and NI PCIe-8381) with an IMEC PXIe card (IMEC PXIE_1000), recorded with SpikeGLX version 20190413-phase3B2 (https://billkarsh.github.io/SpikeGLX/). Craniotomies were performed the day before recording and covered with Kwik-Cast (World Precision Instruments), and mice were allowed to recover overnight. On the day of recording, a 3D-printed piece was placed over the head of the mouse, blocking from the mouse’s sight the experimenter manipulating the probe above the mouse’s head. Probes were coated in dye (DiD, DiI, or DiO; Thermo Fisher Scientific Vybrant Multicolor Cell-Labeling Kit) to visualize probe tracks in histology. Neuropixels probes were lowered using a Thorlabs micromanipulator (PT1-Z8) at 1-10 *μ*/sec. Slower speeds were used for VTA insertions. The probe was inserted 100 *μ* past the target depth, then retracted to the target depth, and allowed to settle for 10-30 minutes prior to starting the recording.

### Optotagging dopamine neurons

Four shank Neuropixels 2.0 probes were used to record from antidromically optotagged VTA dopamine neurons projecting to lNAc (**Fig. 1k-q, Fig. 3h-o**). In the experiments of **Fig. 1**, ChrimsonR was expressed in dopamine neurons by injecting 300 nL of AAV8-hSyn-FLEX-ChrimsonR-tdTomato (UNC Vector Core) into the VTA of *Dat-Cre^+/-^* mice. During the experiment dopamine axons in lNAc were illuminated with red light (635 nm LED, single pulses, 20 ms pulse duration, 40 mW/mm^2^) at the beginning and end of the recording session (40 trials each); these trials were used to identify lNAc-projecting dopamine neurons (see **Definition of optotagged neurons**). In the experiments of **Fig. 3**, ChR2 was expressed in dopamine neurons by injecting 300 nL of AAV5-Ef1a-fDIO-hChR2(H134R)-EYFP (UNC Vector Core) into the VTA of *Dat-Flp^+/-^;Tac1Cre^+/-^*mice. Dopamine axons in lNAc were illuminated with blue light (473 nm laser, LaserGlow, single pulses, 20 ms pulse duration, 80 mW/mm^2^) at the beginning and end of the recording session (60 trials each).

### Optotagging D1-MSNs

Single shank Neuropixels 1.0 probes were used to record from optotagged D1-MSNs in lNAc (**Fig. 3a-g**). ChrimsonR was expressed in D1-MSNs by injecting 300 nL of AAV8-hSyn-FLEX-ChrimsonR-tdTomato (UNC Vector Core) into the lNAc of *Tac1-Cre^+/-^*mice. Optogenetic light pulses (635 nm LED, 40 mW/mm^2^) were 1 ms instead of 5 ms, to separate light artifacts from light-triggered spiking responses, which typically had latency ≈ 2-3 ms (**Extended Data Fig. 6a**). The first light pulse of the opto-stim patterns (**Fig. 3a**; excluding patterns 3 and 4) was used to identify optotagged D1-MSNs (see **Definition of optotagged neurons**).

### Muscimol injections

Muscimol (Sigma-Aldrich M1523) was dissolved at 1ng/nL in sterile saline (0.9% NaCl) and aliquots were frozen at −20°C. Prior to experiments, an aliquot was thawed, further diluted to a final concentration of 0.5ng/nL in saline, and used for several days, stored at 4°C. One day before starting muscimol/saline injections, mice were anesthetized with isoflurane, and a craniotomy was made above lNAc (from bregma: AP: 1.3 mm, ML: 1.8 mm) and covered with KwikCast (World Precision Instruments). Mice were allowed to recover overnight. Injections of muscimol or saline (alternating days) then occurred the next day and every subsequent day for up to 8 days. For injections, mice were head-fixed and awake within the recording setup. The craniotomy was exposed, rinsed with saline, and visualized using a color camera (Chameleon3 CM3-U3-13S2C, FLIR). Muscimol or saline (50-100 nL) was injected into the lNAc (DV: 4.0 mm) using a glass pipette (Drummond, pulled using Narishige PC-10) and oil hydraulic microinjector (Narishige MO-10). After 10 minutes, the pipette was extracted from the brain. Mice then rested in their home cage for 1-2 hours before starting the behavioral task and photometry recording.

### Multisite D1-MSN stimulation with GRAB_DA3m_ photometry

In this experiment (**Fig. 5**), a mirrored angled fiber was used to target DLS (MFC_400/430-0.48_3mm_MF1.25_MA45, Doric Lenses), facing towards the lateral side, to restrict red light illumination to DLS. Flat faced fibers were used to target lNAc and DMS. Mice had either 2 or 3 fibers implanted, targeting lNAc + DLS, or lNAc + DMS + DLS. An earlier cohort of mice had DLS implants which were not useable due to experimenter error (incorrect orientation of mirrored angled fiber); in these mice, only data from lNAc + DMS were analyzed.

### Histology

Following experiments, mice were deeply anesthetized with an overdose of ketamine/medetomidine, exsanguinated with 0.9% phosphate buffered saline (PBS), and transcardially perfused with cold 4% paraformaldehyde (PFA) in PBS. The brain was extracted from the skull and stored in 4% PFA for 24-48 hours at 4°C, after which it was rinsed with PBS, stored in PBS, and cut into 100 *μ* sections on a vibratome (VT1000S, Leica). In some experiments, sections were immunostained for tyrosine hydroxylase (AB152, Sigma Aldrich) to visualize dopamine axons or cell bodies. Sections were mounted onto glass slides with a mounting medium containing 4’,6-diamidino-2-phenylindole (DAPI; Vectashield, Vector Laboratories) and imaged with an automated fluorescence microscope (Axio Scan.Z1, Zeiss) or confocal microscope (LSM 700, Zeiss).

### Data analysis

Unless otherwise noted, data were analyzed with custom code written in MATLAB (version 2024a).

### Statistics

All statistical tests were performed in MATLAB (version 2024a). Unless otherwise noted, *P* values reflect two-tailed tests. We did not perform tests for normality or correct for multiple comparisons.

### Data exclusion

In the muscimol experiment (**Fig. 4, Extended Data Fig. 7, Extended Data Fig. 8**), we excluded sessions in which injection (saline or muscimol) visibly failed (no movement of meniscus in glass pipette, possibly due to clogging). In total, 4/30 sessions were excluded (1 saline, 3 muscimol), leaving 26 sessions from 6 mice (16 saline, 10 muscimol). Including these sessions did not change the main pattern of results.

### Photometry preprocessing

Raw photometry signals (F) were converted to deltaF/F (dF/F) through the formula 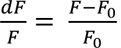, where *F*_0_was defined as the 10^th^ percentile of F in a rolling 30 second window. dF/F traces were upsampled from 20 Hz to 1000 Hz through linear interpolation (MATLAB function *interp1*) and then smoothed with a Gaussian filter (SD = 50 ms). Smoothed dF/F traces were z-scored by subtracting the mean and dividing by the standard deviation of signals in the ITI periods, where ITI periods were defined by excluding time points from 0.5 seconds before to 3 seconds after the start of each trial. GCaMP signals (**Fig. 2, Fig. 4**) were corrected by regressing out the isosbestic signal, using linear regression (function *robustfit* in MATLAB) fit to ITI periods. GRAB_DA3m_ signals were not corrected by the isosbestic channel as the isosbestic wavelength for GRAB_DA3m_ is unknown, and using an incorrect isosbestic wavelength is more likely to distort the signal than correct it (motion artifacts were addressed for these animals using GFP control mice, e.g. **Extended Data Fig. 1c** and **Extended Data Fig. 6g**). Photometry data were synchronized to behavioral data using TTL sync pulses delivered to the photometry recording setup at the start of each behavioral trial.

### Spike sorting

Neuropixels data were spike-sorted offline with Kilosort3^96^ using the ecephys pipeline (Allen Institute, adapted by Jennifer Colonell for SpikeGLX data; https://github.com/jenniferColonell/ecephys_spike_sorting), which includes CatGT to high-pass filter the raw data and detect and remove artifacts (https://github.com/billkarsh/CatGT). Kilosort clusters were then manually curated in Phy (https://github.com/cortex-lab/phy). After manual curation, quality control (QC) metrics and mean waveforms were computed for each unit using code from the ecephys pipeline. Neuropixels data were synchronized to behavioral data using TTL sync pulses delivered to the SMA input of the IMEC PXIe acquisition module at the beginning of each behavioral trial.

### Peristimulus time histograms

Peristimulus time histograms (PSTHs) of neural activity (spike times or photometry) or behavioral data (motion energy) were computed using a MATLAB toolbox written by HyungGoo Kim (https://github.com/HRKimLab/libkm).

### Definition of optotagged neurons

#### Optotagged dopamine neurons

Dopamine neurons were optotagged antidromically^97^ by stimulating dopamine axons in the striatum while recording spiking activity in the VTA with a 4 shank Neuropixels 2.0 probe (**Fig. 1k,l**). To be defined as optotagged, a unit had to meet the following criteria:

1. Pass a stimulus-associated latency test (SALT)^98^ independently in two sets of 40 (**Fig. 1k-q**) or 60 (**Fig. 3h-n**) optotagging trials (single 20 ms pulses of light, red light for ChrimsonR in **Fig. 1** and blue light for ChR2 in **Fig. 3**) delivered at the beginning and end of the recording session (SALT *P* < 0.01 for both sets of trials). SALT compared spiking histograms from 1 to 20 ms (1 ms bins) following laser onset to a null distribution taken from ITI periods.
2. Meet the following quality control criteria based on the QC metrics output by the ecephys pipeline, as recommended by the Allen Institute (https://allensdk.readthedocs.io/en/latest/_static/examples/nb/ecephys_quality_metrics.html): *isi_viol* < 0.5, *amplitude_cutoff* < 0.1, *presence_ratio* > 0.9. These metrics ensured neurons were well-isolated and present throughout the recording session.
3. Have Pearson’s correlation at least 0.9 between mean waveform computed during light delivery (“light-triggered spikes”) and outside light delivery (“spontaneous spikes”). Waveforms were extracted using the software C_Waves written by Bill Karsh (https://github.com/billkarsh/C_Waves).
4. Pass a collision test, a classic test of antidromic activation^97^. The first 1 ms bin following light onset in which firing rate differed significantly from a shuffle control was identified (z-score to shuffle > 5), which we may call *t*_*resp*_. Trials were identified in which spontaneous spikes occurred between *t*_*resp*_ − 7 ms and *t*_*resp*_− 2 ms. To pass the collision test, these trials had to have 0 spikes in the 1 ms bin at *t*_*resp*_. If no spontaneous spikes occurred between *t*_*resp*_− 7 ms and *t*_*resp*_ − 2 ms, the neuron was allowed to pass the collision test.
5. Have mean firing rate > 0.2 Hz and < 10 Hz.

#### Optotagged D1-MSNs

D1-MSNs were optotagged by illuminating lNAc while recording spiking activity with a Neuropixels 1.0 probe (**Fig. 3a-g**). To be defined as optotagged, a neuron had to pass a stimulus-associated latency test (SALT)^98^ using the first pulse of each of the 7 stimulation patterns, excluding patterns 3 and 4 (SALT *P* < 0.01). Light pulse duration was 1 ms in this experiment; SALT compared spiking histograms from 2 to 6 ms (1 ms bins) following laser onset to a null distribution taken from ITI periods.

### Generating z-scored firing rate traces of single neuron spiking activity

Spike trains of individual neurons were transformed into smoothed z-scored firing rate traces by binning spikes (1 ms bins), smoothing the binned firing rate trace with a Gaussian kernel (SD = 50 ms), and z-scoring the resulting trace to ITI periods (**Fig. 1o-q, Fig. 2b,c, Extended Data Fig. 2d-f, Fig. 3h-n, Extended Data Fig. 4**). Here ITI was defined by excluding time points from 0.5 seconds before to 3 seconds after the start of each trial; all other time points were labeled “ITI”. Firing rate traces were z-scored by subtracting the mean and dividing by the standard deviation of ITI time points.

### Definition of neurons significantly excited/inhibited by odors during opto-conditioning task

To define neurons as “significantly excited/inhibited” by an odor (CS+ or CS-) (**Fig. 2b,c, Extended Data Fig. 4**), trials were split into 1 second bins, spanning 1 second before odor onset to 5 seconds after odor onset. The number of spikes in each of the 5 1-second bins following odor onset was compared to the number of spikes in the 1-second bin preceding odor onset using a one-tailed ranksum test, separately for each odor (*n* = number of trials). Any neuron with *P <* 0.001 in any of these 5 comparisons was deemed “significantly excited” (right tail; **Fig. 2b,c, Extended Data Fig. 4a-d**) or “significantly inhibited” (left tail; **Extended Data Fig. 4e**) by the odor.

### Behavioral analysis

#### Video processing

Motion energy was extracted from within ROIs in each video. ROIs were drawn manually around the whisker pad, nose, mouth, and eye for face videos using custom MATLAB code. For body videos, the ROI was defined as the whole frame (full body motion). Within each ROI, motion energy was computed as the absolute value of the difference between subsequent frames.

### Computational modeling

#### Linear time-invariant (LTI) systems identification analysis

Consider an input-output system ℱ, with input denoted by *u*(*t*) and output denoted by *y*(*t*), such that *y*(*t*) = ℱ(*u*(*t*)). This system is called linear time-invariant (LTI) if it has two properties:

1. Linearity:

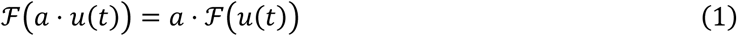

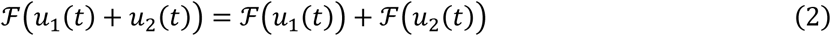
2. Time invariance:

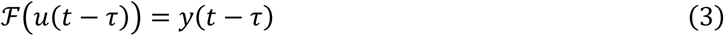

A system is LTI if and only if it is completely characterized by its *impulse response*, ℎ(*t*), i.e. the response of the system to a Dirac delta function input:

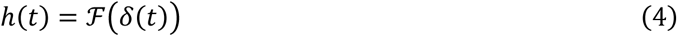

Where *δ*(*t*) is the Dirac delta function (not to be confused with the TD error, which we denote *TD*(*t*)). This means that (with zero initial condition), the output of an LTI system is equal to the convolution of the input with the impulse response (convolution denoted as ∗):

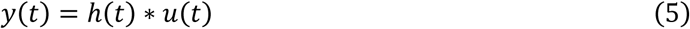

By taking the Laplace transform of both sides of equation 5, we can turn the convolution into a product (the *convolution theorem* of Laplace transforms):

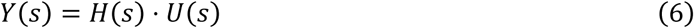

*H*(*s*), known as the *transfer function* of the system, is the Laplace transform of the system’s impulse response, and conversely, the impulse response ℎ(*t*) is the inverse Laplace transform of the transfer function:

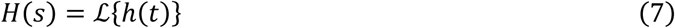

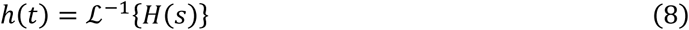

It is useful to consider the subset of all possible transfer functions *H*(*s*) which can be expressed as the ratio of two polynomials with real coefficients *K*, *a_i_*, and *b*_*j*_, with the numerator having order *m* and the denominator having order *n*:

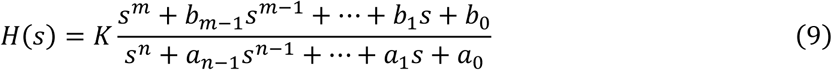

This ratio-of-polynomials form can be factored as follows (fundamental theorem of algebra):

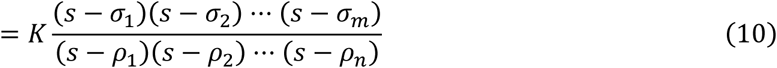

Where these roots, *σ*_i_ and *ρ*_*j*_, are complex numbers and are known as the “zeros” and “poles” of the system, respectively. As will be seen, the zeros and poles are very useful for interpreting and understanding the system dynamics. Many real-world LTI systems have *H*(*s*) with this form, but some do not (e.g. a strict time delay between input and output cannot be expressed this way).

When *H*(*s*) has this form, then the system is equivalent to a linear differential equation with constant coefficients, as follows:

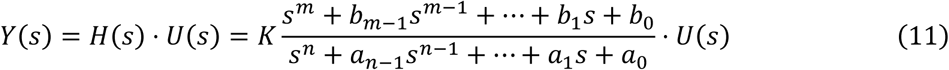

Rearranging terms:

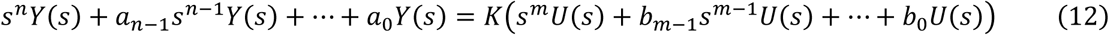

Taking the inverse Laplace transform of both sides:

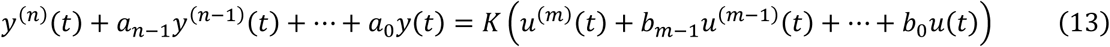

Where the last step follows from the derivative property of Laplace transforms (ℒ{*f*^′^(*t*)} = *s* ⋅ ℒ{*f*(*t*)}). Transforming the system into this form is useful for comparing it to the TD equation, which in the absence of rewards is a linear differential equation with *m* = 1 (1 zero), *n* = 0 (0 poles), K = 1, and *b*_0_ = *σ*_1_ = ln *γ* (see **The TD equation is an LTI system in the absence of rewards**).

Real biological systems are neither linear (e.g. neurons have threshold non-linearities) nor time-invariant (e.g. brain states like arousal fluctuate over seconds to minutes). We wished to understand the extent to which the transformation from striatal D1-MSN spiking to midbrain dopamine activity (spiking or release) can be described as an LTI system (i.e., how good of an approximation is this?), and to find the best fit ℎ(*t*) describing this transformation (what is its shape? How many zeros and poles are needed to describe the system?). Finally, we wished to relate this empirical ℎ(*t*) to the TD equation, which is itself an LTI system in the absence of rewards, transforming the value function *V*(*t*) into the TD error *TD*(*t*) by convolving with the TD impulse response, ℎ_*TD*_ (*t*) (see **The TD equation is an LTI system in the absence of rewards).**

#### The TD equation is an (improper) LTI system in the absence of rewards

The continuous form of the TD equation is^21^:

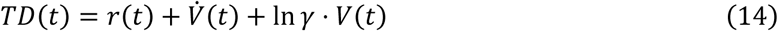

In the absence of rewards (*r*(*t*) = 0), this reduces to:

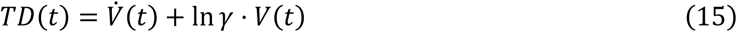

This is an improper (non-causal) LTI system with 0 poles and 1 zero, *σ*_1_ = −ln (*γ*). The TD equation outputs zero when:

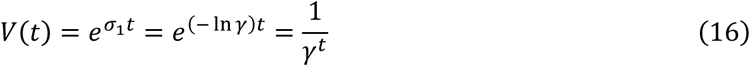

The TD equation with *r*(*t*) = 0 is equivalent to convolving the value function *V*(*t*) with an impulse response function, which we call the “TD impulse response” or ℎ_*TD*_ (*t*):

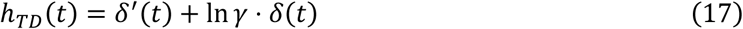

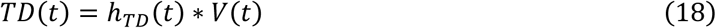

Where *δ*(*t*) is the Dirac delta function and *δ*′(*t*) is the derivative of the Dirac delta function.

#### Fitting LTI models to data

To fit LTI models to data (**Fig. 3d-n, Fig. 5c-e, Extended Data Fig. 6e-m, Extended Data Fig. 9**), we used MATLAB’s System Identification Toolbox, specifically the function *tfest*. LTI models were fit to the average PSTH for each stimulation pattern. To fit an order *n* model we used *tfest* with *n* zeros and *n* + 1 poles (e.g. **Fig. 3f**). No regularization was used. Cross-validated R^2^ was computed by fitting the model on 6 of 7 stimulation patterns and predicting the response to the held-out pattern; these held-out predictions were concatenated across the 7 patterns, and cross-validated R^2^ was computed as 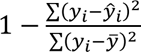, where *y* is the true value of the average PSTH at time-point *i*, 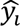 is the prediction of the LTI model, and *y̅* is the mean of all *y_i_*. To fit to a derivative model (order 1 model with *σ*_1_ fixed at 0), we first calculated the derivative of the input (MATLAB function *diff*), forcing *σ*_1_ = 0, and then fit an LTI model with 0 zeros and 2 poles. The cross-validated R^2^ of this model was compared against that of a full order 1 LTI model with 1 zero and 2 poles (**Fig. 3g,k, Extended Data Fig. j-l**).

#### Estimating *γ* from bootstrapped dopamine neuron population activity

We estimated *γ* from bootstrapped dopamine neuron population activity as follows. First, we sampled 55 optotagged dopamine neurons randomly with replacement from the full set of 55 neurons. Then we computed the average PSTH of this bootstrapped population and re-fit the LTI model (1 zero, 2 poles) predicting the average PSTH from the stimulation patterns. The estimated gamma, 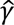, was computed using the formula 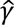= *e*^−*σ*^^1^, where *σ*_1_ is the zero of the LTI fit (see above for derivation). This process was repeated 1000 times to generate the histogram in **Fig. 3m**. For each bootstrap, we also computed the ratio P/N (the ratio of the positive and negative areas of the fit impulse response). We plotted P/N against 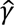 across all 1000 bootstraps in **Fig. 3o**. The black line in **Fig. 3o** shows an exponential curve of the form *ae^bx^* + *c*, fit to the data using the MATLAB function *fit*.

## Acknowledgements

We thank HyungGoo Kim for help setting up the antidromic optotagging technique; Ryunosuke Amo for technical advice; Edward Soucy, Brett Graham, and Yuwei Li of the Harvard Neurotechnology Core for engineering assistance; the Harvard Center for Biological Imaging for imaging support; Harvard FAS Research Computing for computing support; Bernardo Sabatini, Samuel Gershman, and Peter Dayan for critical feedback; and all Uchida lab members for their input. This work was supported by grants from the National Institute of Health (NIH) (5U19NS113201 to N.U. and 5R01DA05975 to N.U. and M.W.-U.) and the Simons Collaboration on Global Brain (to N.U.). M.G.C. is supported by a NIH K99/R00 Pathway to Independence Award (1K99DA060290).

## Author contributions

M.G.C. and N.U. designed the experiments. M.G.C., Y.R., Z.C., and S.X. collected data. S.M. assisted with establishing experimental procedures. M.G.C., Z.C., and M.B. developed the LTI analysis framework. M.G.C. analyzed the data. M.G.C. wrote the first draft of the manuscript and created the figures. All authors edited the manuscript.

## Data and code availability

Data and code will be posted to online repositories upon publication.

## References

1. Schultz, W., Dayan, P. & Montague, P. R. A neural substrate of prediction and reward. Science 275, 1593–1599 (1997).

2. Bayer, H. M. & Glimcher, P. W. Midbrain dopamine neurons encode a quantitative reward prediction error signal. Neuron 47, 129–141 (2005).

3. Steinberg, E. E. et al. A causal link between prediction errors, dopamine neurons and learning. Nat. Neurosci. 16, 966–973 (2013).

4. Amo, R. et al. A gradual temporal shift of dopamine responses mirrors the progression of temporal difference error in machine learning. Nat. Neurosci. 1–11 (2022).

5. Kim, H. R. et al. A Unified Framework for Dopamine Signals across Timescales. Cell 183, 1600–1616.e25 (2020).

6. Howhy, J. The Predictive Mind. (Oxford University Press, Oxford, England, 2013).

7. Sutton, R. S. Learning to predict by the methods of temporal differences. Mach. Learn. 3, 9–44 (1988).

8. Sutton, R. S. & Barto, A. G. Reinforcement Learning, Second Edition: An Introduction. (MIT Press, 2018).

9. Arulkumaran, K., Deisenroth, M. P., Brundage, M. & Bharath, A. A. Deep reinforcement learning: A brief survey. IEEE Signal Process. Mag. 34, 26–38 (2017).

10. Lillicrap, T. P., et al. Continuous control with deep reinforcement learning. arXiv [cs.LG] (2015).

11. Mnih, V. et al. Human-level control through deep reinforcement learning. Nature 518, 529–533 (2015).

12. Ainslie, G. Specious reward: a behavioral theory of impulsiveness and impulse control. Psychol. Bull. 82, 463–496 (1975).

13. Frederick, S., Loewenstein, G. & O’donoghue, T. Time discounting and time preference: A critical review. J. Econ. Lit. 40, 351–401 (2002).

14. Mischel, W., Shoda, Y. & Rodriguez, M. I. Delay of gratification in children. Science 244, 933–938 (1989).

15. Jeong, H. et al. Mesolimbic dopamine release conveys causal associations. Science 378, eabq6740 (2022).

16. Coddington, L. T., Lindo, S. E. & Dudman, J. T. Mesolimbic dopamine adapts the rate of learning from action. Nature 614, 294–302 (2023).

17. Hamid, A. A. et al. Mesolimbic dopamine signals the value of work. Nat. Neurosci. 19, 117–126 (2016).

18. Qian, L. et al. Prospective contingency explains behavior and dopamine signals during associative learning. Nat. Neurosci. 1–13 (2025).

19. Howe, M. W., Tierney, P. L., Sandberg, S. G., Phillips, P. E. M. & Graybiel, A. M. Prolonged dopamine signalling in striatum signals proximity and value of distant rewards. Nature 500, 575–579 (2013).

20. Mohebi, A. et al. Dissociable dopamine dynamics for learning and motivation. Nature 570, 65–70 (2019).

21. Mikhael, J. G., Kim, H. R., Uchida, N. & Gershman, S. J. The role of state uncertainty in the dynamics of dopamine. Curr. Biol. 32, 1077–1087.e9 (2022).

22. Gershman, S. Dopamine ramps are a consequence of reward prediction errors. Neural Comput. 26, 467–471 (2014).

23. Chang, C. Y. et al. Brief optogenetic inhibition of dopamine neurons mimics endogenous negative reward prediction errors. Nat. Neurosci. 19, 111–116 (2016).

24. Maes, E. J. P. et al. Causal evidence supporting the proposal that dopamine transients function as temporal difference prediction errors. Nat. Neurosci. 23, 176–178 (2020).

25. Houk, J. C., Davis, J. L. & Beiser, D. G. Models of Information Processing in the Basal Ganglia. (MIT Press, 1995).

26. Kawato, M. & Samejima, K. Efficient reinforcement learning: computational theories, neuroscience and robotics. Curr. Opin. Neurobiol. 17, 205–212 (2007).

27. Keiflin, R. & Janak, P. H. Dopamine Prediction Errors in Reward Learning and Addiction: From Theory to Neural Circuitry. Neuron 88, 247–263 (2015).

28. Parker, N. F. et al. Choice-selective sequences dominate in cortical relative to thalamic inputs to NAc to support reinforcement learning. Cell Rep. 39, 110756 (2022).

29. Cromwell, H. C. & Schultz, W. Effects of expectations for different reward magnitudes on neuronal activity in primate striatum. J. Neurophysiol. 89, 2823–2838 (2003).

30. Samejima, K., Ueda, Y., Doya, K. & Kimura, M. Representation of action-specific reward values in the striatum. Science 310, 1337–1340 (2005).

31. Ito, M. & Doya, K. Distinct neural representation in the dorsolateral, dorsomedial, and ventral parts of the striatum during fixed- and free-choice tasks. J. Neurosci. 35, 3499–3514 (2015).

32. Lowet, A. S. et al. An opponent striatal circuit for distributional reinforcement learning. Nature 639, 717–726 (2025).

33. Cox, J. & Witten, I. B. Striatal circuits for reward learning and decision-making. Nat. Rev. Neurosci. 20, 482–494 (2019).

34. Lammel, S. et al. Input-specific control of reward and aversion in the ventral tegmental area. Nature 491, 212–217 (2012).

35. Menegas, W., Akiti, K., Amo, R., Uchida, N. & Watabe-Uchida, M. Dopamine neurons projecting to the posterior striatum reinforce avoidance of threatening stimuli. Nat. Neurosci. 21, 1421–1430 (2018).

36. Saunders, B. T., Richard, J. M., Margolis, E. B. & Janak, P. H. Dopamine neurons create Pavlovian conditioned stimuli with circuit-defined motivational properties. Nat. Neurosci. 21, 1072–1083 (2018).

37. Farassat, N. et al. In vivo functional diversity of midbrain dopamine neurons within identified axonal projections. Elife 8, (2019).

38. de Jong, J. W. et al. A Neural Circuit Mechanism for Encoding Aversive Stimuli in the Mesolimbic Dopamine System. Neuron 101, 133–151.e7 (2019).

39. de Jong, J. W., Liang, Y., Verharen, J. P. H., Fraser, K. M. & Lammel, S. State and rate-of-change encoding in parallel mesoaccumbal dopamine pathways. Nat. Neurosci. 27, 309–318 (2024).

40. Klapoetke, N. C. et al. Independent optical excitation of distinct neural populations. Nat. Methods 11, 338–346 (2014).

41. Zhuo, Y. et al. Improved green and red GRAB sensors for monitoring dopaminergic activity in vivo. Nat. Methods 1–12 (2023).

42. Coddington, L. T. & Dudman, J. T. In Vivo Optogenetics with Stimulus Calibration. in Patch Clamp Electrophysiology: Methods and Protocols (eds. Dallas, M. & Bell, D.) 273–283 (Springer US, New York, NY, 2021).

43. Oettl, L.-L. et al. Phasic dopamine reinforces distinct striatal stimulus encoding in the olfactory tubercle driving dopaminergic reward prediction. Nat. Commun. 11, 3460 (2020).

44. Yagishita, S. et al. A critical time window for dopamine actions on the structural plasticity of dendritic spines. Science 345, 1616–1619 (2014).

45. Iino, Y. et al. Dopamine D2 receptors in discrimination learning and spine enlargement. Nature 579, 555–560 (2020).

46. Lee, S. J. et al. Cell-type-specific asynchronous modulation of PKA by dopamine in learning. Nature 590, 451–456 (2021).

47. Long, C. et al. Constraints on the subsecond modulation of striatal dynamics by physiological dopamine signaling. Nat. Neurosci. 27, 1977–1986 (2024).

48. Masset, P. et al. Multi-timescale reinforcement learning in the brain. Nature 642, 682–690 (2025).

49. Sousa, M. et al. A multidimensional distributional map of future reward in dopamine neurons. Nature (2025) doi:10.1038/s41586-025-09089-6.

50. Kobayashi, S. & Schultz, W. Influence of reward delays on responses of dopamine neurons. J. Neurosci. 28, 7837–7846 (2008).

51. Ljung, L. System Identification: Theory for the User, Second Edition. (Prentice Hall, 1998).

52. Kramer, P. F., Twedell, E. L., Shin, J. H., Zhang, R. & Khaliq, Z. M. Axonal mechanisms mediating γ-aminobutyric acid receptor type A (GABA-A) inhibition of striatal dopamine release. Elife 9, (2020).

53. Howe, M. W. & Dombeck, D. A. Rapid signalling in distinct dopaminergic axons during locomotion and reward. Nature 535, 505–510 (2016).

54. Coddington, L. T. & Dudman, J. T. The timing of action determines reward prediction signals in identified midbrain dopamine neurons. Nat. Neurosci. 21, 1563–1573 (2018).

55. Engelhard, B. et al. Specialized coding of sensory, motor and cognitive variables in VTA dopamine neurons. Nature 570, 509–513 (2019).

56. Watabe-Uchida, M. & Uchida, N. Multiple Dopamine Systems: Weal and Woe of Dopamine. Cold Spring Harb. Symp. Quant. Biol. 83, 83–95 (2018).

57. Tsutsui-Kimura, I. et al. Distinct temporal difference error signals in dopamine axons in three regions of the striatum in a decision-making task. Elife 9, (2020).

58. Verharen, J. P. H., Zhu, Y. & Lammel, S. Aversion hot spots in the dopamine system. Curr. Opin. Neurobiol. 64, 46–52 (2020).

59. Bogacz, R. Dopamine role in learning and action inference. Elife 9, (2020).

60. Akiti, K. et al. Striatal dopamine explains novelty-induced behavioral dynamics and individual variability in threat prediction. Neuron 110, 3789–3804.e9 (2022).

61. Lee, R. S., Sagiv, Y., Engelhard, B., Witten, I. B. & Daw, N. D. A feature-specific prediction error model explains dopaminergic heterogeneity. Nat. Neurosci. 27, 1574–1586 (2024).

62. Mohebi, A., Wei, W., Pelattini, L., Kim, K. & Berke, J. D. Dopamine transients follow a striatal gradient of reward time horizons. Nat. Neurosci. 27, 737–746 (2024).

63. Calipari, E. S., Huggins, K. N., Mathews, T. A. & Jones, S. R. Conserved dorsal-ventral gradient of dopamine release and uptake rate in mice, rats and rhesus macaques. Neurochem. Int. 61, 986–991 (2012).

64. Critchfield, T. S. & Kollins, S. H. Temporal discounting: basic research and the analysis of socially important behavior. J. Appl. Behav. Anal. 34, 101–122 (2001).

65. Time and Decision: Economic and Psychological Perspectives of Intertemporal Choice. (Russell Sage Foundation Publications, 2003). doi:10.7758/9781610443661.

66. Story, G. W., Vlaev, I., Seymour, B., Darzi, A. & Dolan, R. J. Does temporal discounting explain unhealthy behavior? A systematic review and reinforcement learning perspective. Front. Behav. Neurosci. 8, 76 (2014).

67. Matousek, J., Havranek, T. & Irsova, Z. Individual discount rates: a meta-analysis of experimental evidence. Exp. Econ. 25, 318–358 (2022).

68. Vuchinich, R. E. & Simpson, C. A. Hyperbolic temporal discounting in social drinkers and problem drinkers. Exp. Clin. Psychopharmacol. 6, 292–305 (1998).

69. Enomoto, K., Matsumoto, N., Inokawa, H., Kimura, M. & Yamada, H. Topographic distinction in long-term value signals between presumed dopamine neurons and presumed striatal projection neurons in behaving monkeys. Sci. Rep. 10, 8912 (2020).

70. Ainslie, G. & Haslam, N. Hyperbolic discounting. Choice over time 5, 57–92 (1992).

71. Laibson, D. Golden eggs and hyperbolic discounting. Quarterly Journal of Economics 112, 443–478 (1997).

72. Hayden, B. Y. Time discounting and time preference in animals: A critical review. Psychon. Bull. Rev. 23, 39–53 (2016).

73. Kurth-Nelson, Z. & Redish, A. D. Temporal-difference reinforcement learning with distributed representations. PLoS One 4, e7362 (2009).

74. Peters, A. J., Fabre, J. M. J., Steinmetz, N. A., Harris, K. D. & Carandini, M. Striatal activity topographically reflects cortical activity. Nature (2021) doi:10.1038/s41586-020-03166-8.

75. Tsutsui-Kimura, I. et al. Dopamine in the tail of the striatum facilitates avoidance in threat-reward conflicts. Nat. Neurosci. 28, 795–810 (2025).

76. Greenstreet, F. et al. Dopaminergic action prediction errors serve as a value-free teaching signal. Nature 1–10 (2025).

77. Lutas, A. et al. State-specific gating of salient cues by midbrain dopaminergic input to basal amygdala. Nat. Neurosci. 22, 1820–1833 (2019).

78. Costa, V. D., Dal Monte, O., Lucas, D. R., Murray, E. A. & Averbeck, B. B. Amygdala and Ventral Striatum Make Distinct Contributions to Reinforcement Learning. Neuron 92, 505–517 (2016).

79. Liu, C. et al. An action potential initiation mechanism in distal axons for the control of dopamine release. Science 375, 1378–1385 (2022).

80. Harkin, E. F. et al. Temporal derivative computation in the dorsal raphe network revealed by an experimentally driven augmented integrate-and-fire modeling framework. Elife 12, (2023).

81. Nagel, K. I. & Wilson, R. I. Biophysical mechanisms underlying olfactory receptor neuron dynamics. Nat. Neurosci. 14, 208–216 (2011).

82. Liss, B. & Roeper, J. 3.4 Ion channels and regulation of dopamine neuron activity. Dopamine handbook 118, (2010).

83. Stetsenko, A. & Koos, T. Neuronal implementation of the temporal difference learning algorithm in the midbrain dopaminergic system. Proc. Natl. Acad. Sci. U. S. A. 120, e2309015120 (2023).

84. Abbott, L. F., Varela, J. A., Sen, K. & Nelson, S. B. Synaptic depression and cortical gain control. Science 275, 220–224 (1997).

85. Eshel, N. et al. Arithmetic and local circuitry underlying dopamine prediction errors. Nature 525, 243–246 (2015).

86. Watabe-Uchida, M., Eshel, N. & Uchida, N. Neural Circuitry of Reward Prediction Error. Annu. Rev. Neurosci. 40, 373–394 (2017).

87. Liu, Z. et al. A distinct D1-MSN subpopulation down-regulates dopamine to promote negative emotional state. Cell Res. 32, 139–156 (2022).

88. Grace, A. A. & Bunney, B. S. Induction of depolarization block in midbrain dopamine neurons by repeated administration of haloperidol: analysis using in vivo intracellular recording. J. Pharmacol. Exp. Ther. 238, 1092–1100 (1986).

89. Tucker, K. R., Huertas, M. A., Horn, J. P., Canavier, C. C. & Levitan, E. S. Pacemaker rate and depolarization block in nigral dopamine neurons: a somatic sodium channel balancing act. J. Neurosci. 32, 14519–14531 (2012).

90. Groves, P. M., Wilson, C. J., Young, S. J. & Rebec, G. V. Self-inhibition by dopaminergic neurons. Science 190, 522–528 (1975).

91. Yang, H. et al. Nucleus Accumbens Subnuclei Regulate Motivated Behavior via Direct Inhibition and Disinhibition of VTA Dopamine Subpopulations. Neuron 97, 434–449.e4 (2018).

92. Simpson, E. H. et al. Lights, fiber, action! A primer on in vivo fiber photometry. Neuron 112, 718– 739 (2024).

93. Cohen, J. Y., Haesler, S., Vong, L., Lowell, B. B. & Uchida, N. Neuron-type-specific signals for reward and punishment in the ventral tegmental area. Nature 482, 85–88 (2012).

94. Jun, J. J. et al. Fully integrated silicon probes for high-density recording of neural activity. Nature 551, 232–236 (2017).

95. Steinmetz, N. A. et al. Neuropixels 2.0: A miniaturized high-density probe for stable, long-term brain recordings. Science 372, eabf4588 (2021).

96. Pachitariu, M., Sridhar, S. & Stringer, C. Solving the spike sorting problem with Kilosort. bioRxiv 2023.01.07.523036 (2023) doi:10.1101/2023.01.07.523036.

97. Lipski, J. Antidromic activation of neurones as an analytic tool in the study of the central nervous system. J. Neurosci. Methods 4, 1–32 (1981).

98. Kvitsiani, D. et al. Distinct behavioural and network correlates of two interneuron types in prefrontal cortex. Nature 498, 363–366 (2013).

